# The Effect of Compositional Context on Synthetic Gene Networks

**DOI:** 10.1101/083329

**Authors:** Enoch Yeung, Aaron J. Dy, Kyle B. Martin, Andrew H. Ng, Domitilla Del Vecchio, James L. Beck, James J. Collins, Richard M. Murray

## Abstract

It is well known that synthetic gene expression is highly sensitive to how comprising genetic elements (promoter structure, spacing regions between promoter and coding sequences, ribosome binding sites, etc.) are spatially configured. An important topic that has received far less attention is how the physical layout of entire genes within a synthetic gene network affects their individual expression levels. In this paper we show, both quantitatively and qualitatively, that compositional context can significantly alter expression levels in synthetic gene networks. We also show that these compositional context effects are pervasive both at the transcriptional and translational level. Further, we demonstrate that key characteristics of gene induction, such as ultra-sensitivity and dynamic range, are heavily dependent on compositional context. We postulate that supercoiling can be used to explain these interference effects and validate this hypothesis through modeling and a series of *in vitro* supercoiling relaxation experiments. On the whole, these results suggest that compositional context introduces feedback in synthetic gene networks. As an illustrative example, we show that a design strategy incorporating compositional context effects can improve threshold detection and memory properties of the toggle switch.

## INTRODUCTION

A fundamental aspect of designing synthetic gene networks is the spatial arrangement and composition of individual genes. With advancements in DNA assembly technology (Engler et al., 2008; Gibson et al., 2009; Lee et al., 2015; Weber et al., 2011), drops in sequencing and prototyping costs (Chappell et al., 2013; Shin & Noireaux, 2012; Sun et al., 2013), the continual discovery of novel synthetic biological parts (Stanton et al., 2014), synthetic biology is poised to make a quantum leap in the size and complexity of the networks it can build. So why hasn’t it happened yet?

The challenge is that synthetic biological parts can be highly sensitive to context (Cardinale & Arkin, 2012), e.g. the physical composition of elements in synthetic genes, conditions of the host chassis, and environmental parameters. Context effects can often be mitigated by engineering principles such as standardization (Davis et al., 2011; Mutalik et al., 2013a) or high-gain feedback (Del Vecchio et al., 2008; Mishra et al., 2014). Frequently, it is critical to have an understanding of physical mechanisms underlying context effects before they can be resolved (Davis et al., 2011; Mutalik et al., 2013a). The key insight is that context effects in synthetic gene networks can rarely be ignored; the study of context effects leads to principle-based design approaches that mitigate their interference.

Complementary to principle-based design approaches are large-library screening approaches (Kosuri et al., 2013). Kosuri et al. showed that it is possible to rapidly screen combinatorial promoter-ribosome-coding sequence libraries for intended gene expression levels and regulatory function, even if models for individual genetic elements such as promoters and RBSs have limited prediction power (Kosuri et al., 2013). Smanski et al. (2014) screened a large combinatorial library for a sixteen gene nitrogen fixation cluster, to explore the effect of genetic permutations in ordering, orientation, and operon occupancy. They discovered there were strong differences in nitrogenase activity, depending on the compositional configuration, but no clear architectural trends emerged from monitoring acetylene reduction. Moreover, the number of compositional context variants (more than *𝒪*(10^19^)) of a sixteen gene cluster made it impossible to exhaustively search and screen for the optimal variant.

These results underscore the complementary role that library screening and principle-based design approaches have in synthetic biology. Library screening approaches can be an extremely effective way to optimize performance in individual parts. However, the number of possible compositional context variants for larger biological networks quickly mushrooms to scales that are intractable for library-based approaches. If we are to build increasingly larger synthetic biocircuits, including synthetic genomes designed from scratch (Gibson et al., 2010), we need a deeper physical understanding of how compositional context affects gene expression.

Most recently, Chong and coworkers showed that transient accumulation of localized positive supercoiling can lead to reduction in gene expression — they showed through *in vitro* transcription experiments that supercoiling could be a physical mechanism behind transcriptional bursting (Chong et al., 2014). Their results also suggested that the presence of nearby topological barriers such as DNA-bound proteins or transcriptional activity of neighboring genes can affect local gene expression.

To paraphrase John Donne, the broad implication of these studies is that “no [gene] is an island entire of itself". Indeed, Rhee et al. (1999); Shearwin et al. (2005) show that genes with overlapping transcripts are subject to transcriptional interference. However, even in non-overlapping genes, the statistical analysis of Korbel et al. (2004) suggest there is a strong link between spatial arrangement and co-regulated gene pairs. Is the same true of synthetic gene networks? Has the field of synthetic biology ignored a fundamental mode of transcriptional regulation in natural gene networks that could be exploited in synthetic gene networks? How does compositional context, i.e. the spatial arrangement of genes, affect gene expression in synthetic gene networks?

## RESULTS

### Compositional context significantly affects transcription of synthetic genes

To explore compositional context, we constructed a set of plasmids, varying gene orientation, relative orientation, coding sequence identity, and the length of spacing between genes. There are three relative orientations that two genes can assume: 1) convergent orientation, where transcription of both genes proceeds in opposite directions and towards each other, 2) divergent orientation, where transcription of both genes proceeds in opposite directions, away from each other and towards genetic elements on the plasmid backbone, and 3) tandem orientation, where transcription of both genes proceeds in the same direction (Liu & Wang, 1987; Shearwin et al., 2005). We constructed plasmids of each orientation to examine their effect on gene expression *in vivo* and *in vitro.*

Each plasmid incorporated two reporter genes, assembled and inserted in the same locus of a consistent vector backbone. Each gene consisted of an inducible promoter, the Lac or Tet promoter, and encoded expression of a fluorescent reporter. Each plasmid (containing an orientation variant) was transformed into MG1655Z1 *E. coli*, which expresses LacI and TetR constitutively from the genome. Since LacI and TetR do not cross-regulate each other, we were able to conclude that any expression differences across strains were solely due to context effects (Ceroni et al., 2015).

We first used mSpinach RNA aptamer and MG RNA aptamer as reporters downstream of the Lac and Tet promoter, respectively. Since mSpinach RNA aptamer is not cytotoxic, it can be used in live-cell imaging to explore how induction response of the Lac promoter varies with compositional context. In our experiments, we first trapped single cells in a microfluidic chamber (CellASIC ONIX B04A system). Next, we flowed 3,5-difluoro-4-hydroxybenzylidene imidazolinone (DFHBI) in culture media for 30 minutes, the dye that fluoresces when bound to mSpinach, to ascertain background levels of fluorescence from DFHBI. Then we flowed 1 mM of ispropyl-*β*-D-1-thiogalactopyroside (IPTG) to release repression of LacI, thus activating expression of mSpinach RNA aptamer. We observed that the induction response of the Lac promoter varied significantly depending on its relative gene orientation, even though the neighboring gene was never activated by aTc (Figure 2).

**Figure 1:**
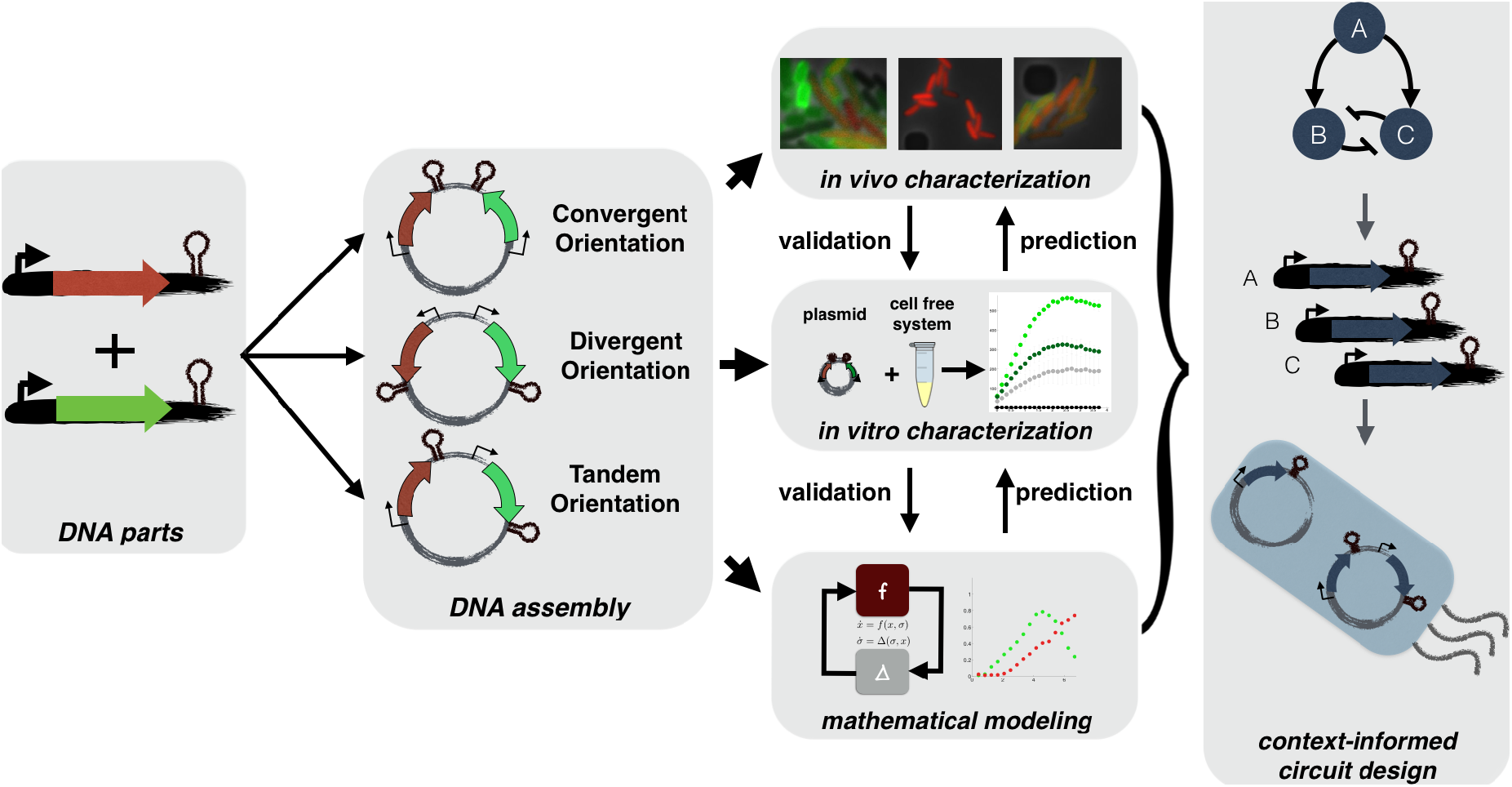
Experimental and theoretical approaches for understanding compositional context effects: Plasmids are constructed using synthetic biology techniques varying gene orientation, transcript length, coding sequence, replication origin, and antibiotic marker. Each plasmid is characterized thoroughly *in vivo* based on what is appropriate for the fluorescent reporter, e.g. single cell fluorescence microscopy, flow cytometry or plate reader. Plasmids are tested *in vitro* in a cell-free expression system to infer physical hypotheses driving compositional context effects and compared against models capturing the hypotheses inferred from *in vitro* and *in vivo* data. These hypotheses support a conceptual framework for designing synthetic biocircuits that utilize compositional context interactions.

**Figure 2:**
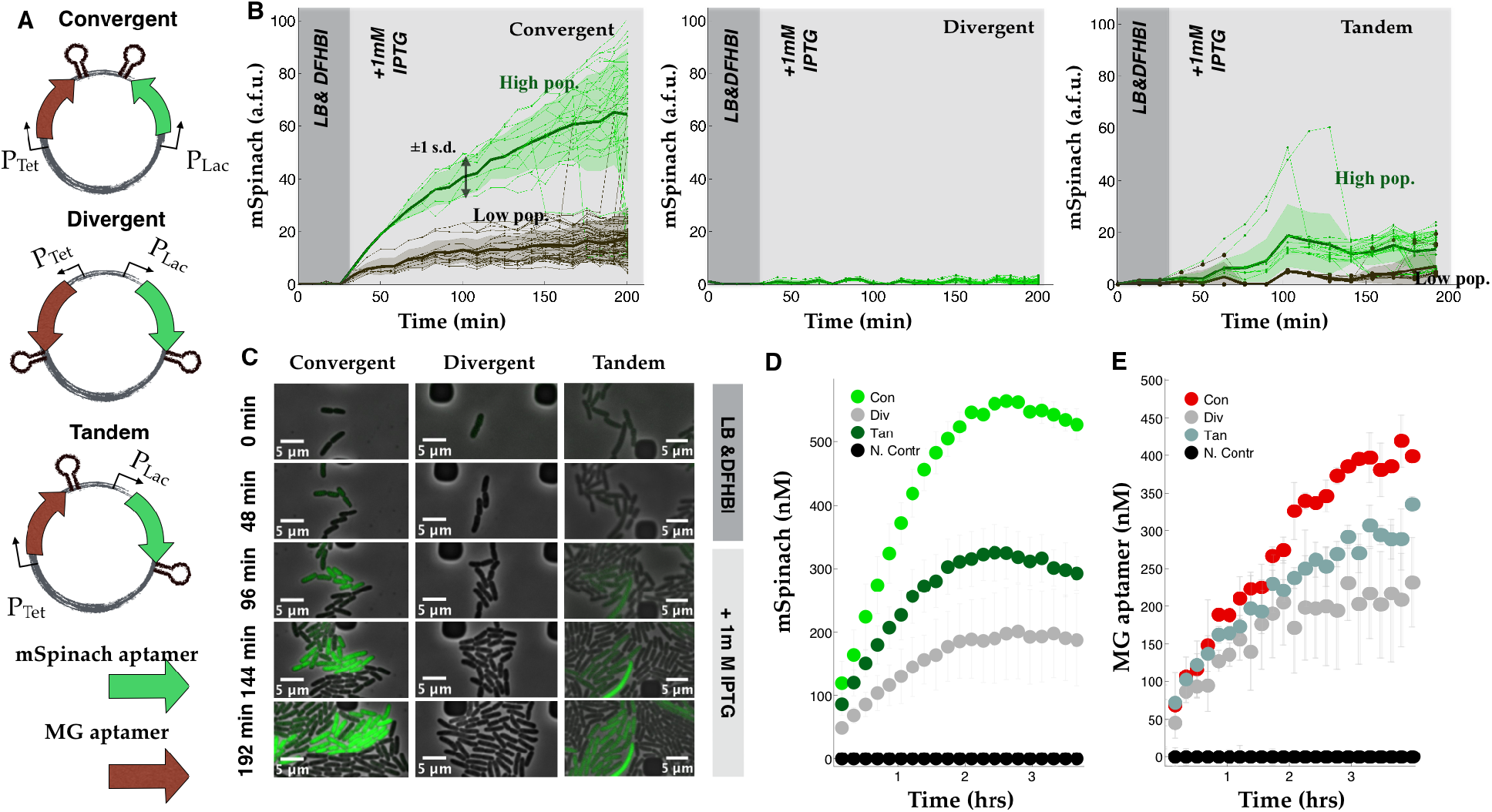
Compositional context alters single cell RNA expression profiles: **(A)** Convergent-, divergent-, and tandem- oriented mSpinach and MG aptamer reporters on ColE1 backbone. **(B)** Time-lapse mSpinach expression curves for individual cell traces in response to 1 mM IPTG induction. Solid central lines within a shaded region denote the mean expression across cell lineages within a population, while the shaded area shows one standard deviation from the mean. **(C)** Single cell microscopy images of convergent-, divergent-, and tandem- oriented mSpinach expression. **(D-E)** Convergent-, divergent-, and tandem- oriented mSpinach and MG RNA aptamer expression in an *E. coli* cell free expression system.

Convergent oriented mSpinach expression produced a ramp-like response to IPTG induction, rising gradually over the course of three hours to reach a steady-state level of expression coinciding with saturation in the microfluidic chamber (see Figure 2). Convergent oriented mSpinach also gave a strong bimodal response to IPTG induction, with one group of cells with high expression (green traces in Figure 2B) and another with low expression (black traces in Figure 2B).

In contrast, divergent oriented mSpinach had a very weak induction response. Tandem oriented mSpinach had bimodal expression as well, with its brightest population of cells expressing at steady-state levels comparable to the weak population in convergent orientation. The remainder of tandem oriented mSpinach *E. coli* cells showed very weak levels of fluorescence.

Interestingly, cells with tandem oriented mSpinach exhibited pulsatile expression, in contrast to the ramp-like response shown by convergent oriented mSpinach. A few outlier cell traces achieved levels of mSpinach expression comparable to the bright convergent mSpinach population, but only at the peak of their transient pulses. Overall, tandem oriented mSpinach exhibited bursty and weaker gene expression than convergent oriented mSpinach.

Since many intracellular parameters fluctuate stochastically *in vivo* (Elowitz et al., 2002), we ran control experiments of each plasmid in a cell-free *E. coli* derived expression system (Noireaux et al., 2003). In this system the effects of single-cell variability are eliminated, e.g. variations in LacI and TetR repressor concentration, polymerase, ribosome, tRNA pools. Also, all deoxynucleotide triphosphates are removed during preparation of cell-extract, thus eliminating any confounding effects of plasmid replication. We prepared separate cell-free reactions for each orientation, assaying mSpinach and MG aptamer expression in a plate reader, using equimolar concentrations for each reaction (Figure 2D-E). Because all cell-free reactions were derived from a single batch of well-mixed extract, the variability in LacI repressor concentration was minimal.

Again, we observed that mSpinach was brightest in the convergent orientation, weakest in the divergent orientation and achieved intermediate expression in the tandem orientation, confirming the trends observed *in vivo.* Likewise, MG aptamer expressed strongest in the convergent orientation, weaker in the tandem orientation, and weakest in the divergent orientation. These *in vitro* outcomes were all consistent with the data from *in vivo* single cell experiments. Since the only connection between our *in vitro* tests and and *in vivo* strains is the plasmids themselves, this conclusively confirms that compositional context is the reason for differences in gene expression. All in all, we conclude that compositional context can significantly alter the transcriptional response of a gene to induction.

### Compositional context effects are pervasive in translational reporters

To explore if these compositional context effects were seen in translated reporters, we first replaced the coding sequence of MG aptamer with the coding sequence for red fluorescent protein (Bba E1010 (Zhang et al., 2002)). We also interchanged the spacer between mSpinach and RFP, to see if our results were dependent on the sequence of the spacer. We then ran an identical experiment, as in Figure 2, to see how induction of mSpinach affected and correlated with RFP expression in single cells.

As expected, relative gene orientation had the same effect on mSpinach transcription as in Figure 2. Even with RFP in place of MG aptamer, mSpinach expression was highest in the convergent orientation and weakest in the divergent orientation.

We also observed that both convergent and tandem oriented mSpinach expressed with a bimodal phenotype (see Supplemental Figure S2B-C). These results confirmed that the identity of the neighboring gene and spacer sequence content was not the source of these gene expression differences.

Interestingly, RFP expression was extremely leaky in the divergent orientation, suggesting that leaky RFP expression somehow broadly interfered with mSpinach expression across all cells. In contrast, convergent oriented mSpinach and RFP showed strong XOR logic — any cells that expressed small amounts of RFP did not respond to IPTG induction with mSpinach expression, while cells that did not express any RFP showed strong mSpinach expression. This data suggests compositional context can be exploited to shape co-expression of neighboring genes.

To further show that compositional context effects extend to translated reporters, we replaced mSpinach with cyan fluorescent protein (CFP) (Veening et al., 2004). We deliberately used a weak RBS, BCD16 from (Mutalik et al., 2013a), for CFP and a strong RBS, derived from pET29-b, from the Bgl Brick pBbE5K plasmid (Lee et al., 2011) to ensure that any ribosome competition effects Gyorgy et al. (2015) would be unidirectional (RFP loading on CFP and not vice-versa) (Gyorgy et al., 2015). Thus, if both genes are induced, any differences in RFP expression would elucidate compositional context effects and not competition for translational resources.

First, we ran a single induction experiment, analogous to experiments run for Figure 2 and Supplemental Figure S2. Induction of CFP with IPTG showed that mean CFP expression was again strongest in the convergent, (slightly) weaker in the tandem orientation, and weakest in the divergent orientation (Figure 3, Supplemental Figures S3C-D (IPTG-only condition).

**Figure 3:**
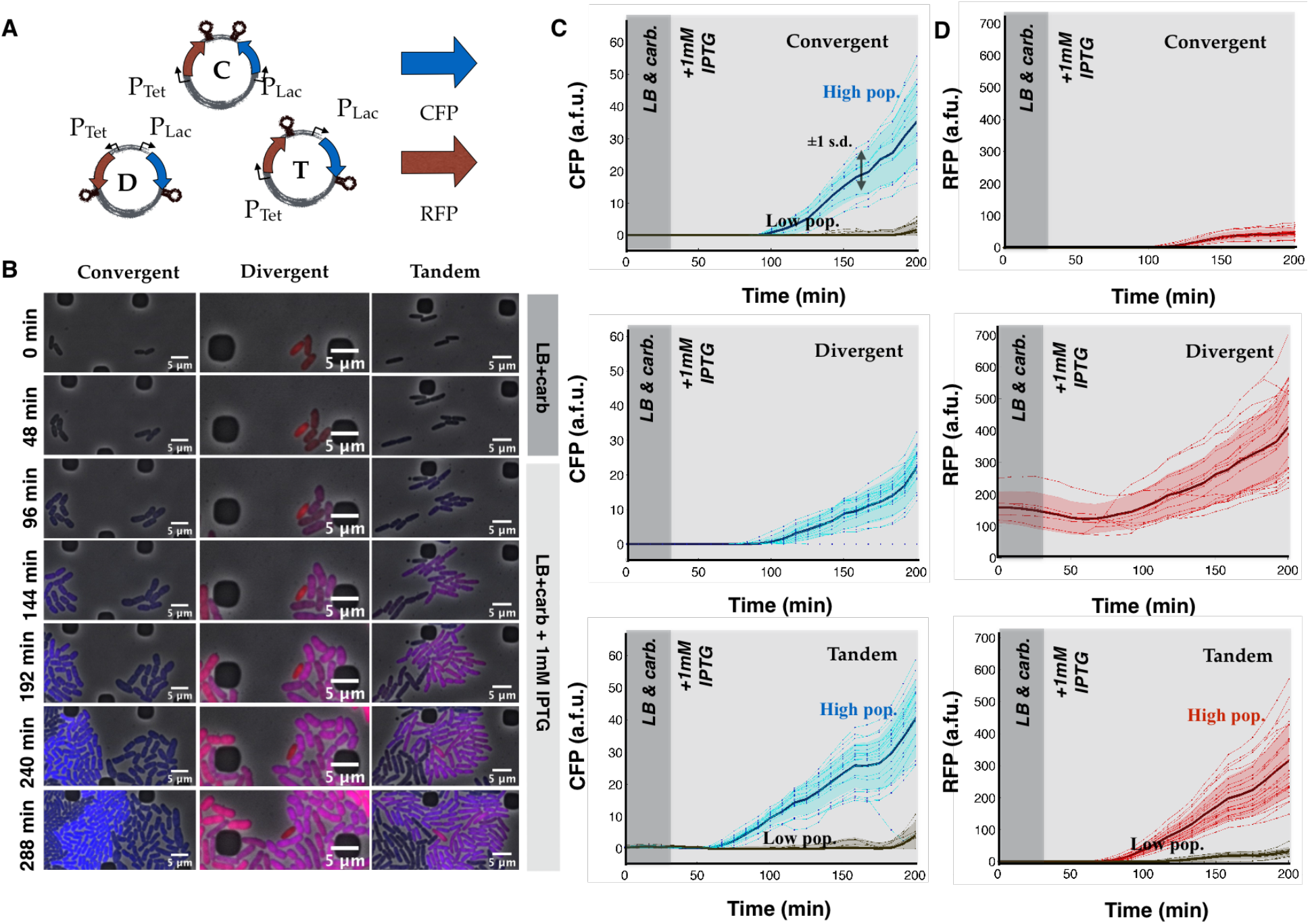
Consistent compositional context effects are observed in translated reporters: **(A)** Plasmid maps for convergent, divergent, and tandem oriented CFP and RFP on the ColE1 plasmid backbone. **(B)** Single cell microscopy images of convergent-, divergent-, and tandem- oriented CFP and RFP expression superimposed. **(C)** Single cell traces of CFP response to IPTG induction in the convergent, divergent, and tandem orientation. Note that both convergent orientation and tandem orientation exhibit bimodal expression phenotypes. High expressing CFP cells also corresponding to high expressing RFP cells, while low expressing CFP cells (gray traces)correspond to low expressing RFP cells. **(D)** Single cell traces of RFP response to IPTG induction in the convergent, divergent, and tandem orientation. Note that both divergent orientation and tandem orientation respond to IPTG induction with significant RFP expression, while convergent orientation responds slightly with some leaky RFP expression.

As a control, we cloned and induced CFP as a single gene on the exact same plasmid locus (either in sense or antisense orientation relative to the plasmid vector). In both control plasmids, 100 bp flanking upstream and downstream sequences were preserved as in the experimental plasmids, to eliminate any promoter sensitivity to upstream sequence perturbation. We noticed a dramatic 5-fold increase in signal over the weakest expressing orientation (compare Supplemental Figures S3B,F-G and S3C). In contrast, comparing sense and anti-sense expression of CFP showed only a small (at most 10% difference in expression). This confirmed that the observed compositional context effects could not be attributed to genetic elements within the plasmid backbone.

We also tested the effect of changing the plasmid vector (from ColE1 to p15A). Plasmid vectors were chosen to feature entirely different compositions of replication origin and resistance marker (Supplemental Figure S4B-C). While the quantitative differences in expression changed by varying plasmid vector (most likely reflecting a change in the copy number of the plasmid), the trends were qualitatively identical. This confirmed that plasmid backbone composition was not the primary source of the observed context effects.

Once again, to control for single-cell variability *in vivo*, we tested RFP and CFP expression of each context variant in a cell free expression system (Shin & Noireaux, 2012). CFP and RFP expressed strongest in convergent orientation, weaker in tandem orientation, and weakest in divergent orientation (Figure 5B). These results were consistent with results of our prior *in vitro* tests with mSpinach-MG aptamer plasmids.

Taken in whole, these findings lead us to conclude that the increase in convergent and tandem CFP expression over divergent oriented CFP was unrelated to resource loading effects, plasmid copy number variability or processes related to plasmid replication. These trends were also consistently observed across multiple coding sequences, transcript lengths, including transcriptional and translational reporters. Therefore, we conclude the compositional context is the primary source of the observed differences in gene expression.

### Induction Response of Genes is Affected Significantly by Compositional Context

To see how compositional context altered the induction response over a range of inducer concentrations, we titrated both IPTG and aTc and quantified RFP and CFP expression in bulk culture plate reader experiments (Figure 4 and Supplemental Figure S3E). As predicted by our choice of RBSs (using a strong RBS for RFP and a weak RBS for CFP), increases in RFP expression (corresponding to increasing aTc concentrations) consistently resulted in decreased CFP expression independent of orientation. As expected, increasing CFP expression did not decrease RFP expression. What was most notable was how gene orientation affected the induction response of RFP expression to varying amounts of aTc inducer.

**Figure 4:**
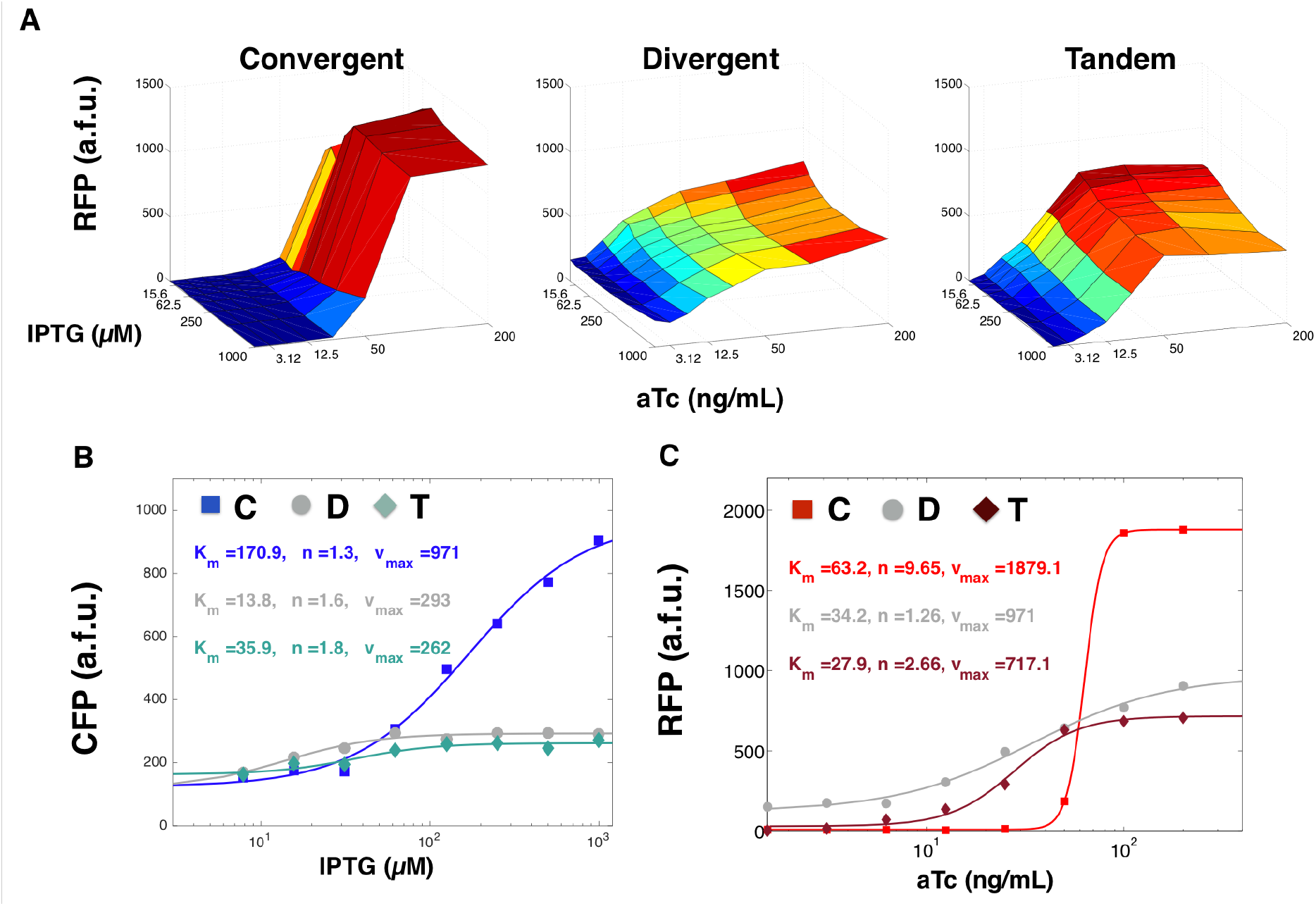
Ultrasensitivity, basal expression, and amplitude of gene induction responses are significantly affected by compositional context: **(A)** RFP expression in a two variable titration assays of IPTG and aTc for convergent, divergent, and tandem oriented CFP RFP plasmids on ColE1 backbone. IPTG is titrated left to right with 2x dilutions starting from 1 mM IPTG (far left) while aTc is titrated top to bottom with 2x dilutions starting from 200 ng/mL. **(B)** Induction curves for convergent-, divergent-, and tandem- oriented CFP fitted to a Hill function at aTc = 200 ng/mL and varying concentrations of IPTG. **(C)** Induction curves for convergent-, divergent-, and tandem- oriented RFP fitted a Hill function at IPTG = 1000 nM and varying concentrations of aTc.

In the convergent orientation, we saw that the transfer curve of RFP expression exhibited strong ultra-sensitivity, increasing by 120-fold in response to only an 8-fold change in aTc. At 100-200 ng/mL of aTc, RFP expression plateaued in an on-state of expression and below 25 ng/mL, RFP expression plateaued in an off-state of expression. Thus, diluting aTc with an 8x dilution factor had the effect of completely switching RFP from an on to an off state.

In contrast, the tandem orientation required a 64-fold change in aTc concentration to achieve a comparable (100x) fold-change in RFP expression. At 100-200 ng/mL, we also saw RFP expression plateaued in an on-state of expression (for all concentrations of IPTG tested). However, RFP reached an off-state of expression only when aTc was diluted down to3 ng/mL or lower. Thus, to achieve the same dynamic range as convergent RFP required an 8x increase in dilution factor.

Divergent oriented RFP exhibited the smallest dynamic range. Varying aTc concentration 200-fold produced at most a 2.7 fold change in RFP expression, despite a 128x dilution factor. Even at low concentrations of aTc, RFP expressed at much higher levels than background. To investigate this effect, we quantified RFP expression in the absence of aTc and discovered it was generally leaky (see Supplemental Figure S3B). We observed similar leaky expression at the single cell level, both in the divergent oriented RFP and CFP MG1655Z1 strain (Figure 3) and divergent oriented RFP and mSpinach strain (Supplemental Figure S3). Since both strains used different spacing sequences of lengths ranging from 150-350 bp, we concluded these leaky effects were a function of RFP gene orientation and not spacer identity nor proximity to the Lac promoter.

We also fit the induction response of each fluorescent protein (titrating the appropriate inducer) while maximally inducing the other gene (Figure 4B-C). Our fits characterized induction response in terms of four parameters, leaky expression *l*, effective cooperativity *n*, maximum expression *V*_*max*_, and half-max induction *K*_*m*_. We noticed that convergent oriented RFP showed significantly increased cooperativity coefficient, nearly four-fold more than tandem orientation and and eight-fold more than divergent orientation. Also, convergent orientation consistently fitted with the highest *K*_*m*_ value in both RFP and CFP induction curves, suggesting that orienting genes convergently raises the induction threshold.

Our experimental data conclusively show that gene expression, induction, and repression can be affected by compositional context. Overall, compositional context can dramatically alter canonical properties of synthetic gene expression and thus should not be overlooked when designing synthetic gene networks.

### A dynamic model incorporating supercoiling states recapitulates observed compositional context effects

Building on the work of (Chong et al., 2014; Liu & Wang, 1987) we investigated whether supercoiling can explain the compositional context effects seen in our data. We constructed an ordinary differential equation (ODE) model describing transcription and translation of both genes. To describe the interplay between gene expression and accumulation of supercoiling for each gene, we introduced separate states to keep track of promoter supercoiling and coding sequence supercoiling. This model structure allowed us to study how supercoiling affected both the processes of transcription initiation and elongation (Drolet, 2006)).

The kinetic rates of transcriptional initiation and transcriptional elongation have been found to be affected significantly by supercoiling buildup (Drolet, 2006). Negatively supercoiled DNA tends to facilitate transcription initiation in the promoter, while transcriptional elongation benefits from negative supercoiling up to a certain point (since negative supercoiling also facilitates melting of the DNA helix). Too much negative supercoiling can lead to the formation of R-loops, structural complexes that involve DNA binding to nascently produced RNA still attached to RNA polymerase. These R-loops complexes have been shown to cause transcriptional stalling (Drolet, 2006).

Conversely, positive supercoiling of DNA introduces torsional stress since positive supercoils naturally oppose the left-handed twist of DNA. Such stress leads to localized regions of tightly wound DNA that is less likely to be transcribed; positive supercoils downstream of a transcription bubble can also impose torsional resistance against further unwinding of the DNA, thereby stalling transcription. When a gene expresses and produces positive supercoiling downstream of the transcription bubble, the accumulation of positive supercoiling can be exacerbated by the presence of a topological barrier, e.g. the binding of a protein such as a transcription factor, or even the presence of another active gene in negatively supercoiled state (Chong et al., 2014). This buildup in positive supercoiling leads to the reduction in the initial rate of gene transcription, or what is often referred to as a bursty profile of gene expression. Thus, excessive twist in the DNA double helix in either direction can decrease transcription rates.

In our model we account for the above considerations by encoding a dependency of transcription rate parameters on local supercoiling density. We build on the analysis of Meyer and Beslon (Meyer et al., 2014) and consider transcription initiation rates to be *dynamically* dependent on supercoiling density. We model them as Hill functions of the absolute deviation of the promoter supercoiling state from a natural super-coiling state (Rhee et al., 1999). In other words, as DNA becomes too twisted in either the positive or negative direction, transcription initiation rates and elongation rates decrease. Similarly, we suppose that the elongation rate of the gene of interest (mSpinach aptamer, MG aptamer, CFP, or RFP) can be modeled as a Hill function of the supercoiling state over the transcript region. Thus, by modeling the dependency of transcription rates on supercoiling, we effectively introduce context-specific coupling between neighboring genes (Figure 5).

**Figure 5:**
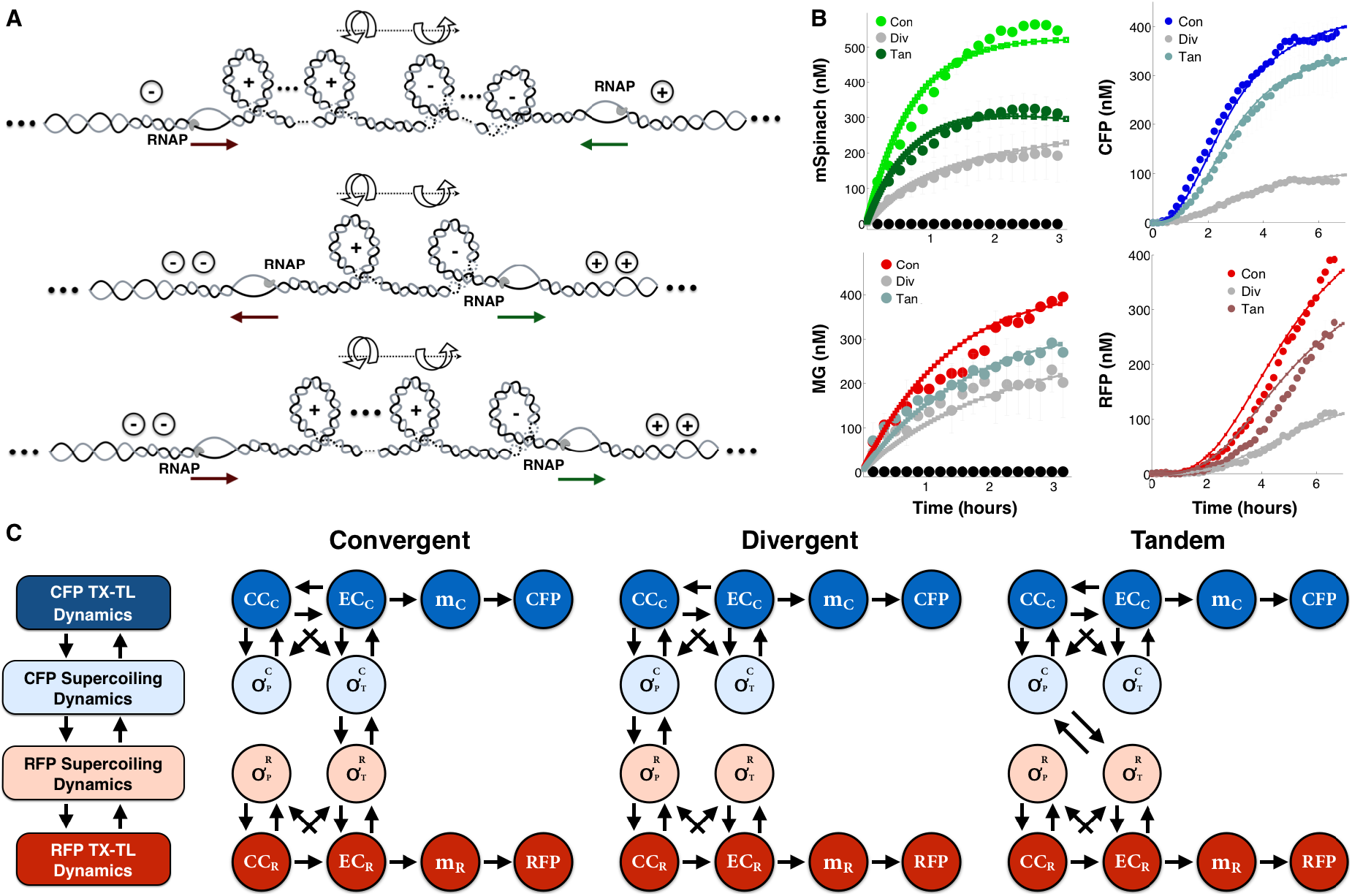
Mathematical models incorporating supercoiling dynamics are able to recapitulate experimental data: **(A)** A diagram showing how positive supercoiling builds up downstream of transcription bubbles and negative supercoiling builds up upstream of transcription bubbles. When two genes are adjacently placed, the intermediate region is exposed to opposing forces of torsional stress from positive (left-handed twist) and negative (right-handed twist) supercoiling. These forces do not cancel out each other, but rather oppose each other to achieve a dynamic equilibrium dependent on the transcriptional activity of nearby genes. **(B)** Expression curves of a mathematical model, integrating supercoiling dynamics of promoter and transcript states with gene expression, with supercoiling parameters fit to experimental data from the TX-TL cell free expression system (Shin & Noireaux, 2012). **(C)** Schematic illustrating the state-dependencies between traditional transcriptional states and supercoiling states in convergent-, divergent-, and tandem- oriented RFP and CFP.

After incorporating these supercoiling hypotheses, our model was able to recapitulate compositional context trends observed in our experimental data (Figure 5B). Our simulations showed that convergent oriented mSpinach (and CFP) is able to achieve higher levels of expression than its divergent and tandem counterparts. Since mSpinach expresses in the anti-sense direction, its expression introduces negative-supercoils upstream according to the twin-domain model by Liu & Wang (1987). In moderate amounts, negative supercoiling facilitates the unwinding of DNA and thus enhances the amount of transcriptional initiation and elongation occurring over the Lac promoter and mSpinach transcript. In the divergent and tandem orientation, mSpinach is expressed in the sense direction, which results in positive supercoiling buildup downstream of the promoter (Figure 5A). The outcome is that divergent and tandem mSpinach expression is reduced (compared to convergent), since the buildup of positive supercoiling inhibits initiation and elongation. This effect is more severe in the divergent orientation, since excessive positive and negative supercoiling generated by initiation of the Tet and Lac promoter can interfere with each other’s expression (Figure 5C).

In exploring the parameter space of our model, we also found that gyrase (an enzyme that relaxes positive supercoiling) and TopoI (an enzyme that relaxes negative supercoiling) activity are not sufficiently high to counteract the coils introduced by rapid repeated transcription events on DNA with two genes. These findings were consistent with the analysis of Chong et al. (2014), Liu & Wang (1987), and Meyer et al. (2014), which argued that buildup of transcription-induced supercoiling far outpaces the activity of supercoiling maintenance enzymes in *E. coli.* This explains why we are able to see compositional context effects both *in vivo* and *in vitro* where gyrase and topoisomerase enzymes are presumably present and active. These results also suggested that extended pre-incubation of plasmids with gyrase would allow us to infer the effect of relaxing positive supercoiling on gene expression in each orientation.

### Relaxing positive supercoiling in plasmids significantly reduces compositional context effects

To test the effect of incubating context-variant plasmids with gyrase, we purified plasmids expressing convergent, divergent, and tandem oriented RFP and CFP from uninduced MG1655Z1 *E. coli.* We divided each plasmid sample into two aliquots — one aliquot was used as a control for the absence of gyrase treatment and the second aliquot was incubated with gyrase (NEB) at 37° C overnight. All gyrase treated samples were prepared and concentrated to equimolar concentrations, so that the enzyme-substrate ratio was identical across all samples. After gyrase incubation, plasmids were purified again to wash out the gyrase buffer using Amicon 0.5 mL Ultracentrifugal Filters. Once again, we tested the expression levels of each plasmid in the cell-free TX-TL system Shin & Noireaux (2012).

In the absence of gyrase, we see that the expression differences between different orientations are largest when comparing convergent and divergent CFP and RFP expression levels. Convergent oriented plasmids expressed almost 300 nM CFP and 500 nM RFP more than its divergent counterpart (see Figure 6). Similarly, tandem oriented plasmid expressed 200 nM more CFP and 300 nM more RFP than divergent orientation. Convergent plasmid also expressed higher than tandem plasmid in both RFP and CFP channels.

**Figure 6:**
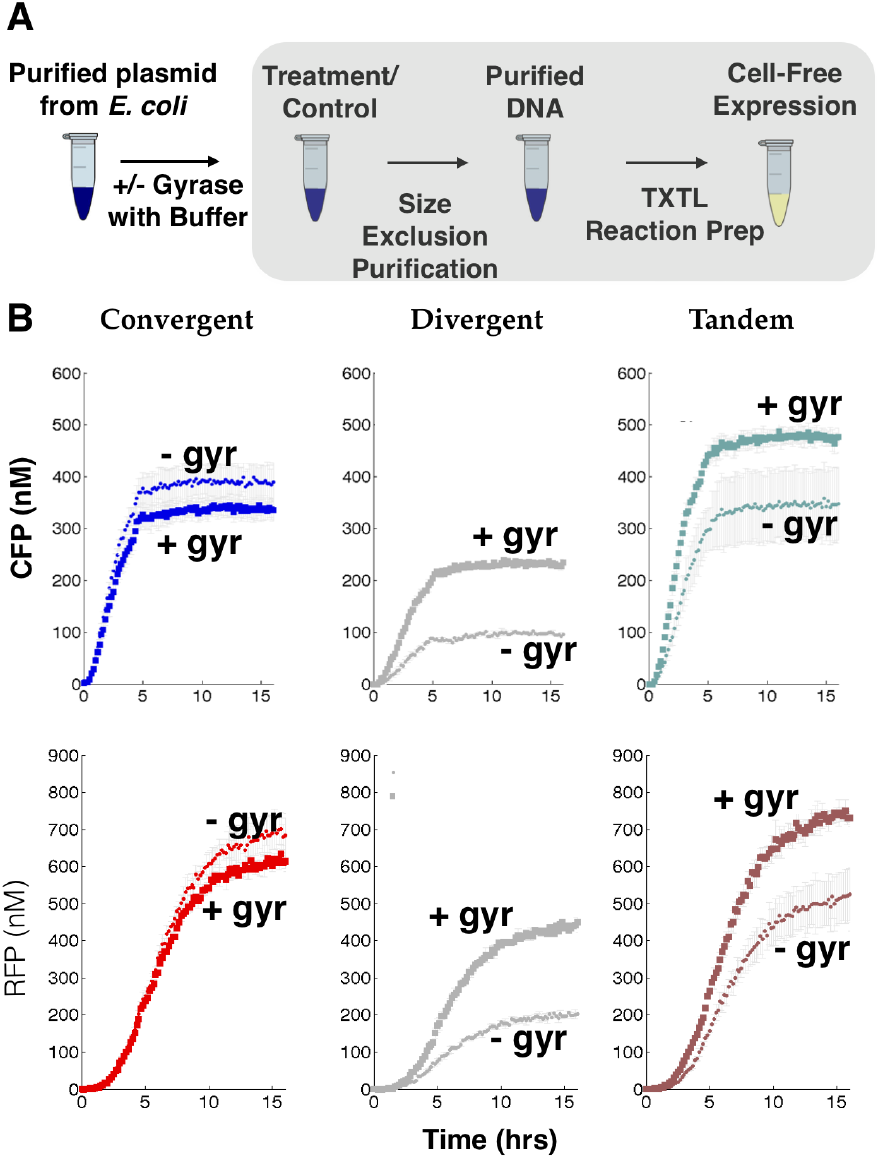
Relaxation of positive supercoiling in plasmids with gyrase enzyme significantly reduces compositional context effects on gene expression: **(A)** Workflow for gyrase treatment experiments. **(B)** Expression of CFP and RFP for convergent, divergent, and tandem oriented ColE1 plasmids prior (small dots) and post treatment (large dots) with gyrase.

After gyrase treatment, tandem oriented CFP and RFP expressed brighter than their convergent and divergent counterparts. We also saw that the disparity in protein expression between convergent and divergent orientation shrunk to 100 nM (66% for CFP) and 180 nM (64% for RFP). Since gyrase serves only to relax plasmids of positive supercoiling, this confirmed that supercoiling is the mechanism underlying compositional context effects.

We anticipated that treatment with gyrase would release positively supercoiled domains in the downstream region of tandem oriented CFP and RFP and release positive supercoiling buildup from divergently (leaky) expressed RFP (Figure 5A) and thereby reduce torsional stress in the promoters of divergently oriented CFP and RFP. Our experimental results confirmed these hypotheses, with divergent orientated CFP and RFP increasing by more than 2 fold and tandem orientated CFP and RFP increasing by 1.4 fold.

Interestingly, gyrase treatment of convergent oriented CFP and RFP appeared to reduce signal slightly, by approximately 10%. This may be because convergent oriented CFP and RFP exhibited little or no leak when uninduced (in contrast to divergent and tandem orientation); thus the purified plasmid for convergent orientation did not have as much positive supercoiling for gyrase to mitigate. Treatments with gyrase may actually have introduced too much negative supercoiling, leading to the small drop in expression observed.

These experimental outcomes are consistent with our model of supercoiling and its impact on compositional context. Gyrase relaxes positively supercoiled domains downstream of convergent and tandem oriented RFP, while in the divergent orientation, gyrase relaxes any positive supercoiling buildup near the promoter region. Once these positive super-coils are removed, the genes are able to express at much higher levels than prior to treatment.

Our data shows that compositional context can have a strong effect on the dynamics of supercoiling within plasmids. Nearby *transcriptionally active* genes or protein-bound genes act as topological barriers to stop migration of supercoils or dispersion of localized torsional stress. Protein-bound genes in particular, act to trap supercoiling in neighboring transcriptionally active genes; this may explain why in our IPTG induction experiments, the mere presence of a repressed RFP and MG gene (respectively) could have such a significant effect on CFP and mSpinach expression. In this way, gene orientation and placement can introduce a fundamentally different form of feedback coupling between neighboring genes. When used appropriately, these feedback effects can be beneficial or consistent with the intended architecture of the biocircuit, as we illustrate in the next section.

### Compositional context improves memory and threshold detection in toggle switch

Synthetic gene networks, for the most part, have been designed primarily to avoid one type of compositional context effect: terminator leakage. Terminator leakage can cause positive correlation between a downstream gene with an upstream genes. While this is a noteworthy consideration in designing synthetic gene networks, it is a dramatic oversimplification of what causes compositional context effects.

In contrast, when we incorporate an understanding of how compositional context and supercoiling cause interference in synthetic gene networks, we can actually utilize compositional context to improve or reinforce the feedback architecture of synthetic gene networks.

The original toggle switch provides an excellent case study of how an informed understanding of compositional context can improve design. Being one of the first synthetic bio-circuits ever made, it was constructed in divergent orientation to avoid terminator leakage effects between two mutually repressing genes, LacI and TetR (Gardner et al., 2000; Kobayashi et al., 2004). From the perspective of protein regulation, two proteins provide mutual negative feedback by transcriptional repression. The LacI protein represses TetR production by binding to the Lac promoter upstream of the TetR coding sequence. The TetR protein represses LacI production by binding to the Tet promoter upstream of the LacI coding sequence.

However, we can also build the toggle switch in convergent or tandem orientation. The convergent toggle switch is most appealing, based on several experimental insights: 1) the competing dynamics of positive and negative supercoils between the two genes encodes an additional layer of mutual negative feedback (Figure 7A), 2) the coexpression profiles of RFP and CFP in the convergent orientation (Supplemental Figure S3)E and mSpinach and RFP in the convergent orientation (Supplemental Figure S2) was strongly anti-correlated. All of these properties of compositional context have the potential to enhance or strengthen the existing mutual negative feedback in the toggle switch.

**Figure 7:**
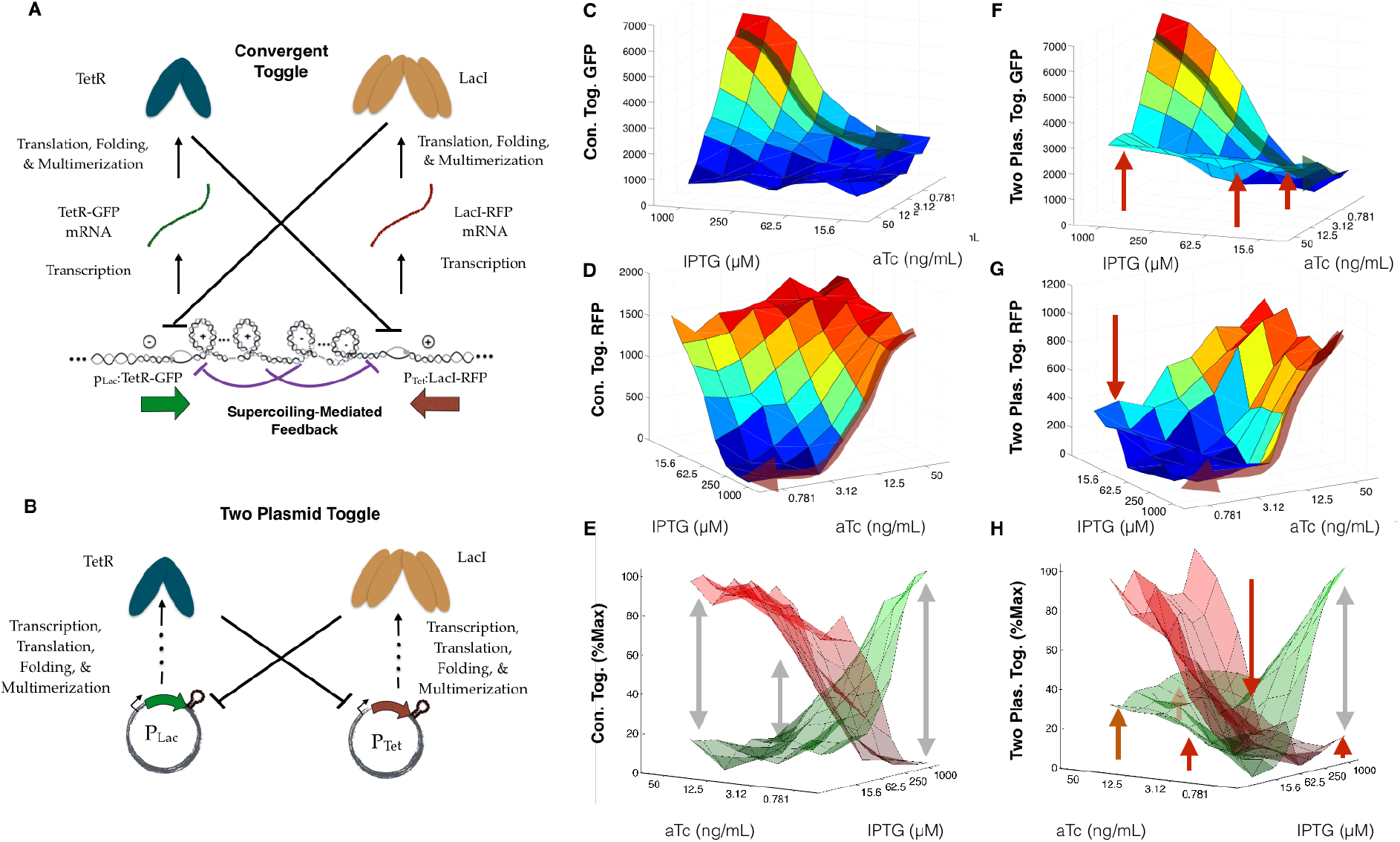
Compositional context can be used to introduce supercoiling-mediated feedback, improving sharpness of threshold in toggle switch: **(A)** Diagram of feedback architecture in a convergent toggle switch. **(B)** Diagram of feedback architecture in a convergent toggle switch TetR-GFP (ColE1) and LacI-RFP (p15A). **(C-E)** Experimental data of convergent toggle GFP expression in response to titrating IPTG and aTc concentration. **(F-H)** Experimental data of the two-plasmid toggle GFP expression in response to titrating IPTG and aTc concentration.

Since our previous controls of sense and anti-sense encoded single genes showed that changing orientation of a single gene on a backbone did not affect expression more than 15 %, we thus constructed a two plasmid version of the toggle switch, with LacI and TetR expressed on separate plasmids. This “context-free" version of the toggle acted as a reference for how a toggle switch should function independent of genetic context.

In both versions of the toggle switch, each gene cassette in the toggle switch was bicistronic, with LacI reported by translation of RFP and TetR reported by GFP. We used stronger ribosome binding sites from the BCD RBS library (Mutalik et al., 2013a), BCD10 and BCD2, to express LacI and TetR and weak BCDs, BCD1 and BCD9, to express the downstream reporters. This was done to minimize any ribosomal loading effects from reporter translation and again, to show that even a toggle switch built *de novo* from existing synthetic biological parts with different CDSs, promoters, and RBSs could utilize compositional context (Figure 7). We also tested the original Gardner-Collins toggle switch, comparing performance in the original orientation to a convergent variant, see Supplemental Figure S4 and discussion in the Supplemental Information.

We first tested the ability of the toggle switch to act as a threshold detector. In theory, the phase portrait of a toggle switch consists of two locally asymptotically stable equilibrium points and a separatrix which drives state trajectories into the basin of attraction of one of the equilibrium points (Gardner et al., 2000). As a proxy for varying the amount of actively repressing TetR and LacI, we simultaneously varied the concentration of inducers aTc and IPTG, thereby allowing us to attenuate the activity of LacI and TetR repression independently. Most notably, when the toggle switch was configured in the convergent orientation, it exhibited much sharper XOR logic and separation between high GFP-low RFP states and high RFP-low GFP states compared to its two-plasmid (and divergent) counterpart (Figure 7 and Supplemental Figure S4A-B).

The two-plasmid toggle exhibits weaker thresholding, i.e. the ability to latch onto a particular state, in two specific parameter regimes — when IPTG and aTc are both present in high concentrations and when IPTG and aTc are both present in low concentrations. When both inducers are present in high concentrations, the majority of Lac and Tet promoters are un-repressed because most repressor proteins are sequestered by inducers, leading to weak feedback. The weak feedback makes it difficult to differentiate which inducer is higher, since essentially all promoters are expressing constitutively (Figure 7F-H). When both inducers are present in low concentrations, both promoters are strongly repressed making it difficult for one promoter to gain a dominant foothold over the other sufficient to produce fluorescent signal. Thus, in the low inducer concentration regime, even if one inducer is higher in concentration than the other, neither gene is strong enough to repress the other to the point of producing detectable fluorescence (Figure 7F-H).

On the other hand, the convergent toggle shows clear separation between high GFP-low RFP states and low GFP-high RFP states in both of these parameter regimes. This improved performance can be explained by examining the effects of supercoiling and compositional context (Figure 7A-B). Suppose, for illustration, that LacI-RFP is slightly more induced than TetR-GFP. The positive supercoiling from TetR-GFP expression propagates downstream to meet the negative supercoils generated from transcription of LacI-mRFP. If LacI-mRFP has propagated negative supercoils, these positive supercoils and negative supercoils create a dynamic equilibrium between opposing torsional forces. As more and more LacI-mRFP expresses, it forces the positive supercoils back into the TetR-GFP coding sequence. When this happens, TetR-GFP is no longer able to express and its transcript region is thus available to LacI-RFP as a downstream region for dissipating its internal torsional stress. Thus, by propagating supercoils into its neighboring gene, LacI-RFP exerts a form of negative feedback independent of transcription factor-mediated repression.

This explains why the convergent toggle is able function in regimes where IPTG and aTc are simultaneously high or low. When IPTG and aTc are both present in high concentrations, the attenuation of transcription factor repression is compensated by the presence of supercoiling mediated repression. Thus, even though LacI and TetR are not as effective in repressing their respective promoters, the extra layer of feedback allows the convergent toggle to decide on a dominant state (LacI-RFP). Similarly, in the low parameter regime, even though both repressors are strong, the additional feedback from supercoiling favors one gene or the other (an enhancement of the winner-takes-all or XOR logic) and evidently improves the ability of the toggle switch to again allow LacI-RFP to dominate over TetR-GFP.

It is worth noting that the supercoiling feedback occurs at a faster time scale (since there is no translation, protein folding, multimerization involved), supporting the usual mode of repression enforced by the LacI transcription factor. The more LacI-RFP expresses, the more torsional stress it exerts on the opposing gene, which strengthens its foothold as the dominant state. Thus, there is a multi-layer feedback effect introduced by supercoiling in the convergent orientation, conformal with the intended feedback architecture of the toggle switch. In this way, we see that compositional context can be a powerful tool for encoding feedback in synthetic gene networks.

## DISCUSSION

### The Link Between Compositional Context Effects and Growth Phase: Temporal Aspects of Compositional Context

Our experimental data, as well the outcomes of several gyrase treatment experiments, support a model of how supercoiling dynamics affect transcription. Depending on its compositional context, the supercoiling state of a gene can be affected by the propagation of supercoiling from nearby coding regions. In this way, supercoiling couples the activity of two neighboring genes. The strength of that coupling and its impact on the temporal dynamics of gene expression depends on the orientation of the genes and what part of the gene is exposed to torsional stress from the neighboring gene. On the whole, these features of our model are able to recapitulate the *in vitro* and *in vivo* trends observed at steady-state, but do not account for aspects of how gyrase and topoisomerase levels are regulated during different growth states.

An interesting facet of these context effects are the temporal dynamics of supercoiled genes, topoisomerase concentrations, and their dependence on cell culture growth phase. Specifically, as *E. coli* cells transition from exponential to stationary phase, plasmid DNA exhibited significantly less negative supercoiled DNA. Balke & Gralla (1987) showed that up to ten negative supercoils could be lost in the pBR322 plasmid in stationary phase cells grown in LB. Thus, gyrase activity (which maintains negative supercoiling) is attenuated as cells approach the end of their exponential growth phase. These findings are corroborated by our data; we also saw that compositional context differences become increasingly dramatic just as cells complete their exponential growth phase (Figure S3C-D).

In this work, we have not made a point to model the temporal dynamics of gyrase and topoisomerase as a function of cellular growth phase since doing so would require rigorous characterizations of gyrase and topoisomerase concentrations through the entire growth cycle. Another interesting extension would be to examine how gyrase dynamics and the compositional context of genes in core metabolic systems affect or modulate the dynamics of metabolism.

### Supercoiling Dynamics Dominate Genetic Context Effects

Our analysis considered supercoiling as the physical basis for generating expression differences. In past work, the primary context effects considered in designing synthetic biocircuits are the effects of terminator leakage and transcriptional interference from overlapping promoter and RBS elements. We claim that supercoiling is the dominant source of compositional context effects observed in our data, justified by the following observations.

First, we see consistent differences in gene expression, even when only one gene is induced and the other remains repressed. If terminator leakage and transcriptional interference were the source of compositional context effects, we would not expect to see any effects in the case of single gene induction. But we do see significant context effects, even only if one gene is induced.

Also, if terminator leakage was the dominant process underlying context effects, transcription of vector coding sequences would cause interference. Thus, in our single reporter control plasmids we would expect to see similar levels of transcriptional interference, especially compared to the divergent orientation (where terminator leakage interference would be minimal from either reporter gene).

However, there is more than a 2-fold difference in expression between divergent expressed CFP and its single reporter counterpart (sense or anti-sense, compare Supplemental Figure S3C with Supplemental Figure 3F-G). The physical presence of a neighboring gene has an effect, even if it is not transcriptionally active. Thus, transcriptional interference via terminator leakage does not explain the data.

Second, if transcriptional interference were the primary driver for context effects, we would expect convergent oriented genes to achieve far weaker levels of expression than divergent or tandem orientation. In theory, transcribing polymerases that managed to leak through two terminators (Larson et al. (2008) characterized the termination efficiency of our terminators at 98%) would collide in the convergent orientation, leading to an increase in abortive transcription events or transcriptional stalling.

Admittedly, we see from our *in vivo* characterizations that early-log phase CFP and RFP expression is weaker in the convergent orientation than the divergent and tandem orientation. The fly in the ointment for this argument is that *both* CFP and RFP expression is *higher* (see Figure 4 and Supplemental Figure S3C-E) in the doubly induced case than in the singly induced case in early log phase, which contradicts the predictions of transcriptional interference theory.

Transcriptional interference also does not account for the sudden rise in convergent oriented expression relative to divergent orientation as cells approach the end of their exponential growth phase. Supercoiling theory, on the other hand, predicts that as gyrase activity wanes, the promoter regions of divergently oriented genes become more positively super-coiled, which inhibits their activity. This positive supercoiling originates from the RFP promoter as it transcribes in the anti-sense direction, thus asymmetrically inhibiting CFP expression and favoring RFP expression (see Figure S3E).

Thirdly, consider the differences in expression *in vitro* of convergent, divergent, and tandem transcribed RFP and CFP (Figure 6). Our experimental characterizations *in vitro* control for variations for plasmid copy number (as a function of orientation), since plasmid replication does not occur in the TX-TL cell-free system (Noireaux et al., 2003). Nonetheless, we see that levels of CFP (and RFP) expression in the convergent and divergent orientation differ by nearly 300 nM (and 500 nM) when purified directly from cells in their natural super-coiled state, whereas treating with gyrase to eliminate positive supercoiling decreases the difference by nearly 70% in both genes. Relaxing positive supercoiling in divergently oriented RFP and CFP with gyrase also allows expression levels comparable to tandem orientation prior to gyrase treatment, while treating tandem oriented RFP and CFP enables expression levels higher than both post-treatment and pre-treatment convergent oriented CFP and RFP. Taken in whole, these observations confirm that the dominant physical process driving the effects of compositional context is supercoiling.

### Compositional Context and DNA Replication

While gene transcription occurs both frequently and rapidly, it is important to consider the effects of plasmid DNA replication on supercoiling. Our *in vitro* cell-free experiments results suggest that there are significant differences in gene expression, depending on whether they are in convergent, divergent, and tandem orientation, both at the transcriptional and translation level. Since plasmid DNA replication does not occur in this system, gene orientation and placement has a significant impact on gene expression in the absence of plasmid DNA replication (see Figures 2 and 5). Moreover, the majority of these *in vitro* trends are qualitatively consistent with the trends observed *in vivo.* Thus, our data suggests that either the trends observed are dominated by the context defined by the two reporter genes (mSpinach, MG, CFP, and RPF) of interest or that plasmid replication only strengthens these trends *in vivo.*

Admittedly, we found a small effect of reversing the direction of a single reporter gene on the ColE1 plasmid backbone, to encode it in either the sense or antisense direction. Comparing the expression levels in Supplemental Figure S3F-G, where *t* = 550 minutes after induction, there is a consistent 15% drop in expression if a sense reporter was expressed in the anti-sense direction. This reduction was observed independent of the choice of CFP and RFP coding sequence or choice of promoter. Examining the growth curves, we see that *t* = 550 minutes is still in log-phase for both induced RFP sense and anti-sense strains. Thus, it appears there is a small effect of orientation on CFP and RFP expression. The context of a single gene in this case is the replication origin and the resistance marker, both of which are approximately separated by 200-250 bp from the reporter gene.

However, when we compare the signal of a single induced sense or anti-sense CFP against their convergent, divergent, or tandem counterparts on the same plasmid backbone at the same point in time (Supplemental Figure S3C), we observe up to a 5.8-fold drop in CFP signal or a 2.4-fold drop in RFP signal even when a repressed adjacent gene is introduced, holding all other genes on the plasmid constant. These results strongly suggest that the primary source of context effects are the neighboring reporter gene rather than plasmid. Plasmid replication may indeed have a small impact on genetic context effects, but the observed trends in this study are dominated by variables of compositional context pertaining to our reporter genes of interest. Since the Lac and Tet promoters used in this study (Lutz & Bujard, 1997) are quite strong compared to average wild-type promoters *E. coli*, it remains the subject of future work to investigate whether plasmid replication dynamics play a larger role in defining genetic context when synthetic promoters strengths are attenuated.

### The Role of Compositional Context in Synthetic Biocircuit Design

Our findings show that compositional context significantly alters gene expression in synthetic gene networks. When appropriately harnessed, compositional context can be used to strengthen or enhance existing feedback loops in the intended biocircuit design. These findings validate prior analysis underscoring the value of accounting for compositional context effects in synthetic biocircuit design (Cardinale & Arkin, 2012).

Broadly speaking, there are many levels of abstraction and ways to define compositional context. Cox et al. investigated how different regulatory elements in existing promoters could be assembled in distal, core, and proximal sites to define a library of new combinatorial promoters (Cox et al., 2007). Similarly, Mutalik et al. showed that the compositional context of a ribosome binding site, specifically sequences downstream of the ribosome binding site could have a significant impact on the effective binding strength of the ribosome (Mutalik et al., 2013a). Using a bicistronic design approach, they showed they were able to better insulate against downstream sequence variability to produce predictable parts. These are examples of the importance of understanding and insulating against *intragenic* compositional context.

The results of our experimental studies emphasize the importance of understanding *intergenic compositional context* effects, i.e. composition of entire genes. We have seen that compositional context effects can cause variations of 3-4 fold of the same gene (promoter, RBS, coding sequence, etc.) simply by rearranging its orientation and the orientation of other neighboring genes. The significance of these outcomes raise an important issue. As *intragenic* context, e.g. choice of BCD, promoter design, polycistronic design, are optimized to produce a functional gene cassette with model-predicted gene expression levels (Kosuri et al., 2013; Mutalik et al., 2013b), how do we ensure these predictions are not confounded by intergenic context as genes are composed?

One solution is to separate genes that need to have precise regulated expression levels on to different plasmids. This strategy has been successful, especially if the goal is not to use context effects to enforce additional layers of feedback. For example, a relaxation based genetic oscillator (Stricker et al., 2008) was developed by keeping the transcriptional units for LacI and AraC on separate plasmids. This oscillator exhibited remarkable robustness, oscillating over a range of experimental conditions and had the beneficial feature of tunability.

However, the drawback of this approach is that separating genes on different plasmids introduces imbalances in gene copy number, which in turn can lead to additional design-build-test cycles to rebalance circuit dynamics. Also, it is often the case that there are too many genes in a biocircuit to isolate individually on separate plasmids. In such settings, the findings of this work are important to consider, as they can be used to inform how to optimally compose adjacent genes.

The effects of adding spacing sequences between genes are complex. Specifically, we varied the amount of spacing between mSpinach and MG aptamer in convergent, divergent, and tandem orientation by adding increments of 100 bps between genes and found that spacing did not have a monotonic effect on decreasing the fold-change across orientations (see Supplemental Figure S1A-C). Most unusual was the sudden drop in signal observed in the divergent and tandem orientation, but not in the convergent orientation with 450 bp of spacing between the two genes. It is possible that since the persistence length of DNA is 150 base pairs, 450 base pairs of spacing facilitates formation of plectonomes (with DNA loops consisting of three 150 bp domains) in the spacing region, which induce torsional stress and inhibit formation or movement of the transcription bubble. The physical basis for these observed trends will be a subject of future research.

In general, genes responded well to induction when induced one by one, though their raw expression levels varied depending on compositional context. This may explain why some circuits in the past have been successfully engineered, with little consideration given to the effects of supercoiling and compositional context. For example, the original toggle switch (oriented divergently) was designed to respond and latch to the presence of a single inducer (Gardner et al., 2000) — it did this well and latched to LacI or TetR dominant states. In contrast, the threshold detection abilities of the toggle were not explored.

Likewise, the fold-change in ’off’ vs ’on’ states of three input and four input AND gates developed by Moon and colleagues (Moon et al., 2012) was strongest when comparing singly induced expression levels against the corresponding fully induced state. Interestingly, the four layer and three logic gates in these biocircuits were compositionally composed so that no two genes involved in any constituent layer of logic were placed adjacent to each other. Pairs of genes involved in logic gates were always separated by an auxiliary backbone gene or placed on separate plasmids. Overall, the success in this work suggest that genes can be insulated by inserting short ‘junk’ transcriptional units in between each other. Engineering approaches for attenuating compositional context effects are a subject of future research.

Finally, we remark that the largest effects from varying pairwise-gene orientation occurs when both genes are induced. However, we noticed that tandem oriented CFP and RFP, mSpinach and RFP, and mSpinach and MG aptamer, responded best to both single and double induction (see Figure 1 and Supplemental Figure S3C-D). Although tandem oriented CFP and RFP experienced roughly a 25% drop in signal compared against their single gene counterparts (Supplemental Figure S4), they respond well to induction when individually induced as well as when induced simultaneously. Additionally, although gyrase treatment of tandem oriented plasmids revealed positive supercoiling build-up, tandem oriented plasmid maintained significant levels of expression prior to treatment, comparable to convergent expression and superior to divergent expression. These findings are also consistent with successfully built biocircuits using tandem orientation, e.g. the repressilator (Elowitz & Leibler, 2000), recently developed 3-node and 5-node novel repressilators(Niederholtmeyer et al., 2015), and an eight gene event detector (data not shown).

## EXPERIMENTAL PROCEDURES

### Plasmid Construction, Assembly, and Strain Curation

Plasmids were designed and constructed using either the Gibson isothermal DNA assembly technique (Gibson et al., 2009) or Golden Gate DNA assembly approach (Engler et al., 2008) using BsaI type II restriction enzyme. All plasmids were cloned into JM109 *E. coli* (Zymo Research T3005) or NEB Turbo *E. coli* (NEB C2984H) strains and sequence verified. Sequence verified plasmids were transformed into MG1655Z1 and MG1655ΔLacI (also lacking TetR) strains of *E. coli.* All plasmids with ColE1 replication origin were transformed and cloned at 29 C to maintain low copy number of the ColE1 replication origin. Sequence verified colonies were grown in LB and the appropriate antibiotic and stored as glycerol stocks (17 % glycerol) at -80° C.

### Single Cell Fluorescence Microscopy

Based on the principles elucidated by Han et al.(Han et al., 2013), we ran all our experiments at 29° C when imaging mSpinach. Cells were revived from glycerol stock overnight at 29° C in LB, diluted to an OD of 0.1 and recovered for 2 hours in log-phase. Cells were then diluted to an density of approximately 5 × 10^6^ cells/mL of LB and loaded into a CellASIC plate. Separate solutions for flowing LB with 200 *µ*M DFHBI and LB with 200 *µ*M DFHBI and 1 mM IPTG were prepared and loaded into reagent wells in the CellASIC ONIX B04A plate for imaging.

Fluorescence and bright field images from time-lapse microscopy were cropped using ImageJ and analyzed in MATLAB with Schnitzcell (Young et al., 2012). For characterizing coexpression of mSpinach and MG RNA aptamer, we used single cell agar pad microscopy, with all cells grown shaking at 29° C in a 96 well plate from overnight recovery until they reached log-phase (~4 hours). Induction occurred by transferring 10 *µ*L of cultures into another 96 well plate into 350 *µ*L of LB with 1mM IPTG and 200 ng/mL aTc.

### Plate Reader Experiments

For plate reader experiments, all cultures were revived from glycerol stock at 37° C in LB and the appropriate antibiotic, followed by redilution to OD 0.05-0.1, recovered at log-phase for 2 hours at 37° C, and then pipetted into a 96 square well glass bottom plate (Brooks Life Sciences MGB096-1-2-LG-L) with the appropriate media, antibiotic and inducer. All measurements were taken on Biotek Synergy H1 plate readers, using the internal monochromomator with excitation (and emission) wavelengths for mSpinach, MG aptamer, CFP, and RFP at 469 nm (and 501 nm) at gain 100, 625 (and 655 nm) at gain 150, CFP at 430 nm (and 470 nm) at gain 61 and 100, RFP at 580 nm (and 610 nm) at gain 61 and 100. For RNA aptamer imaging, all *in vitro* and *in vivo* experiments were performed at 29° C with 200 *µ*M DFHBI (for mSpinach) and 50 *µ*M of Malachite Green dye.

## ACKNOWLEDGMENTS

We thank Ophelia Venturelli for her invaluable inspiration and guidance in this project, Victoria Hsiao, Jin Park, Anu Thubagere, Adam Rosenthal for wonderful ideas on imaging, David Younger, Ania Baetica, Vincent Noireaux, Clarmyra Hayes, and Zachary Z. Sun for guidance and assistance with TX-TL experiments, and Lea Goentoro, Johann Paulsson, Long Cai, Jennifer Brophy, John Doyle, Eric Klavins, and Julius Lucks for insightful conversations.

This work was supported in part by a Charles Lee Powell Foundation Fellowship, a Kanel Foundation Fellowship, a National Science Foundation Graduate Fellowship, National Defense Science and Engineering Graduate Fellowship, Air Force Office of Scientific Research Grant (AFOSR) FA9550-14-1-0060, Defense Threat Reduction Agency Grant HDTRA1-14-1-0006, and Defense Advanced Research Projects Agency Grant HR0011-12-C-0065.

## Supplemental Information: Compositional Context Effects in Synthetic Gene Networks

### Experimental Procedures and Data Analysis

#### Plasmid Assembly

Initial efforts to characterize orientation effects involved cloning plasmids with no spacing DNA between genes. We used Gibson assembly to build these plasmids and naturally found that in the convergent and divergent orientation, the primers used to amplify overhangs had strong secondary structure, which reduced cloning efficiency. Thus, we inserted a minimum of 150 base pairs of randomly generated DNA. DNA sequences were randomly generated in MATLAB, using a custom script and the function `randi()` and subsequently screened for secondary structure in Geneious, a gene designer software. All spacer sequences between genes were determined to have no hairpins or any predicted secondary structure at 37° C before use in cloning workflows.

To construct mSpinach and MG RNA aptamer reporter plasmids, we ordered 500 bp Integrated DNA Technologies gBlocks containing the mSpinach and MG RNA aptamer coding sequences in convergent, divergent, and tandem orientation. Backbones and DNA inserts were amplified and prepared at equimolar concentrations in an isothermal Gibson Assembly, incubated for one hour at 50 C, following the methods of of Gibson et. al (Gibson et al., 2009). Gibson products were subsequently transformed into JM109 Zymogen *E. coli* using a quick-transform protocol, plated at 29° C on LB agar plates with 100 µg/mL of carbencillin. Colonies were screened using standard colony PCR techniques, sequence verified using Operon Sequencing’s overnight sequencing service (both Standard and Power Read products). All strains were sequence verified both in JM109 and experimental strains of MG1655Z1 and MG1655Δ LacI.

To construct mSpinach and RFP plasmids, we used a similar approach as described above, except that we used an RFP coding sequence derived from BglBrick plasmid (pBbE5k-RFP), amplified as a linear double stranded DNA molecule compatible with Gibson assembly. We used an analogous approach to construct CFP and RFP reporter plasmids on the ColE1 backbone. To switch backbones (p15A with chloremphenicol resistance marker) and construct CFP and RFP sense and anti-sense plasmids, we used Golden Gate assembly with BsaI-HF enzyme from New England Biolabs (NEB R3535L). All Golden Gate parts were constructed using an internal protocol with standardized four base pair overhangs. Colonies were screened and sequence verified following the same techniques used for plasmids built by Gibson Assembly. Finally, all plasmids developed for this paper were sequence verified both from Qiagen purified plasmid and as glycerol stock (using Operon’s DNA prep service). Sequence verified strains were stored in 17 % glycerol stocks at -80 C with LB and either 100 *µ*g/mL of carbencillin or 34 *µ*g/mL of chloremphenicol.

#### Imaging RNA aptamers: quantitating mSpinach expression using single cell time-lapse fluorescence microscopy, agar pad microscopy and plate readers

In our preliminary tests, we quickly found that mSpinach RNA aptamer is not particularly bright, compared to GFP, RFP, and other standard fluorescent proteins. Moreover, its brightness depends on the operating temperature of the experiment (Han et al., 2013), since the steady state folding configuration of the mSpinach RNA aptamer depends on temperature. We found that mSpinach signal at 200 *µ*M DFHBI (Lucerna Technologies) was nearly undetectable at 37° C. To minimize photobleaching of mSpinach, we developed a custom Python script to interface with MicroManager (Edelstein et al., 2014), employing the fast shutter of the XFO-citep 120 PC (8 ms resolution) to time exposure of the mSpinach expressing cells to light. To maintain an operating temperature of 29° C we used a custom-built microscopy incubation chamber with a World Precision Instruments Heater controller.

Once the microfluidic plate (EMD Millipore Cell ASIC ONIX B04A) was thermally equilibrated, cells were loaded into the imaging chamber and trapped using a loading protocol provided by EMD Millipore to a density of about 3 cells per field of view. Fluorescence microscopy imaging was performed on an Olympus IX81 inverted fluorescence microscope using a Chroma wtGFP filter cube (450/50 BP excitation filter, 480 LP dichroic beamsplitter, and 510/50 BP emission filter), with an XFO-citep 120 PC light source at 100 % intensity and a Hamamatsu ORCA-03G camera. Following the recommendations of (Han et al., 2013), we limited imaging frequency and exposure to every 10 minutes and for 200 ms, respectively. All experiments were conducted with an untransformed control strain of MG1655Z1 *E. coli* in a parallel microfluidic chamber, to quantify background cell fluorescence in DFHBI. For Figures 2B, 3C-D, and Supplemental Figure S2B we segmented and tracked single cell traces of mSpinach (or RFP and CFP) fluorescence using Schnitzcell (Young et al., 2012) and subtracted background fluorescence from each experimental strain. For each point in time, background fluorescence was defined as the maximum of background chamber fluorescence (quantified using ImageJ as mean fluorescence in a nearby non-occupied area of the microfluidic chamber housing the experimental strain) and background cell fluorescence of the control strain for each frame. The majority of background fluorescence was defined by the background fluorescence in cells, with infrequent fluctuations in background fluorescence due to slight perturbations to the autofocus plane.

We also found that fixing cells with paraformaledehyde lead to inconsistent RNA aptamer fluorescence, with unusually high levels of fluorescence in the negative control (especially in the MG aptamer channel). While fixing cells traditionally allows fixation of protein dynamics, this is not true for imaging mSpinach and MG aptamer. It is possible that fixation alters the permeability of the membrane and enables excessive buildup of MG oxalate dye, which at high concentrations nonspecifically binds to RNA molecules in the fixed cell. For this reason, our experimental technique involved imaging of live cells on agar pads, moving as quickly as possible from agar pad to agar pad, and well after the dynamics of induction had reached steady-state.

In contrast, imaging mSpinach in the cell-free expression system developed by Shin and Noireaux (Shin & Noireaux, 2012) required relatively little effort. Fluorescence quantification was performed on a Synergy H1 Biotek plate reader with 469 nm wavelength excitation, 501 nm wavelength emission. Since TX-TL reactions are typically run at 29 C, this further facilitated formation of the mSpinach RNA aptamer in the 32-2 configuration, see the Supplemental Online Material for Paige et. al. (Paige et al., 2011). We did notice that imaging more frequently than 15 minutes had an effect on the dynamics of mSpinach (presumably due to photobleaching), hence we ran all experiments with 15 minute imaging frequencies.

It is important to note that production of mSpinach in 10 *µ*L bulk volume *in vitro* reactions allows for approximately 10^9^ more copies of mSpinach than produced in a cell, we speculate this greatly increases the detectability of mSpinach signal over *in vivo* assays. We found imaging mSpinach in dense cultures (OD ≈ 1) also produced significant signal above background. Thus, the primary challenges of working with mSpinach is its relatively weak signal per single cell. We anticipate that using the latest version of mSpinach (mSpinach2) or dBroccoli in future tests will greatly improve signal (Filonov et al., 2015).

#### Analysis of Plate Reader Data

To generate the data plotted in Figure 4, Supplemental Figures S1B-C, S3 and S4, we extracted data from the Biotek H1 Synergy plate readers using the Gen5 software package, exported to MATLAB matrices for optical density (OD) and fluorescence intensity in either mSpinach (469 excitation, 501 emission, gain 100), GFP (485 excitation, 525 emission, gain 61) CFP (430 excitation, 470 emission, gain 61) and RFP (580 excitation, 610 emission, gain 61 or 150) channels with inverted (bottom-up) fluorescence acquisition. Each sample was background subtracted, normalized by OD, plotted either as a single time point *t* = 9.2 hours corresponding to the tail-end of exponential growth phase or as complete time traces from *t* = 0 to *t* = 11.7 hours (*t* = 700 minutes). Each strain was grown in duplicate in MatriPlate (Brooks Life Science Systems MGB096-1-2-LG-L) 96 well square well glass bottom plates at 500 *µ* L volumes.

Similarly, for toggle switch data analysis we followed the approach outlined above. We note that given our choice of ribosome binding sites for GFP and RFP (BCD1 and BCD9 respectively), expression of GFP and RFP was weaker to avoid ribosomal loading effects. Thus, we did not see significant signal until 8 hours after initial induction. Signal increased monotonically throughout the experiment, varying depending on the balance of IPTG and aTc induction. Data plotted in Figures 7 were background subtracted and normalized by OD.

To estimate RNA aptamer and protein expression in the TX-TL system, we used data from prior calibration experiments, titrating purified fluorescent protein or RNA aptamer and quantitating expression in the Biotek. Each Biotek was calibrated independently; the results of the calibration were used to back out fluorescent protein from raw AFUs, after background subtraction.

#### Flow Cytometry Experiments and Data Analysis

Flow cytometer experiments were performed using a BD Biosciences Flow Assisted Cell Sorter (FACS) Aria II Flow Cytometer to quantify GFP and RFP fluorescence. GFP fluorescence was detected using a 488 nm laser and 530/30 nm internal bandpass filter while RFP fluorescence was detected using 561 nm laser and a 610/20 nm internal bandpass filter. Each plasmid strain (featuring convergent or divergent orientation of the modified pIKE107 Gardner Collins toggle switch) in either MG1655 *E. coli* or MG1655 Δ LacI *E. coli* was plated on cells from clonal glycerol stocks, grown at 37° C on selective media agar plates overnight. Three colonies were picked from each plate to seed replicate cultures for the experiment. All cell cultures were grown in LB media with carbencillin at 100 *µ*g/mL at 37° C. Cultures were induced with either 50 ng/mL of aTc or 1000 *µ*M of IPTG for 5 hours (defined as the latching period from *t* = −5 to *t* = 0 hours). After latching, cells were diluted with a dilution factor of 1000x, in approximate 8 hour intervals, in selective LB media from *t* = 0 to *t* = 48 hours in the experiment. At t = 0, 24, and 48 hours, cell cultures were rediluted and grown for two hours to reach exponential growth phase and rediluted 1:10 in 1x phosphate buffer saline solution. As a negative control, we quantified GFP, RFP, and CFP fluorescence of an untransformed strain of MG1655 *E. coli* as well as cell-free 1x PBS stock to determine forward and side-scatter gating parameters for background particulate matter.

All flow cytometry data was processed using the FlowJo Software. Cells were gated using an ellipsoidal gate of forward and side-scatter values. We utilized live-gating during data acquisition to obtain approximately 20,000 events. All distributions were plotted as modal percentage vs. GFP intensity (in arbitrary fluorescent units). Modal percentage for a given GFP intensity is defined as the ratio of cell count for the given GFP intensity bin normalized by the cell count for the modal GFP intensity bin, multiplied by 100. This method of plotting eliminates the variability of total counts in sub-populations after gating, while still portraying important features of the distribution such as mode, modal variance and modal kurtosis.

### Supplemental Note on Linear DNA Experiments

Since supercoiling buildup is only possible in certain scenarios, e.g. in the presence of chromosomal binding proteins, topologically constrained plasmids or linear DNA tethered to a scaffold (Chong et al., 2014), we used linear DNA to explore how orientation affects gene transcription in the absence of topological barriers. On linear DNA, divergently oriented mSpinach and MG aptamer have relatively little supercoiling buildup since the linear ends of the DNA enable free rotation of the DNA about the helical axis as transcription occurs.

On tandem oriented DNA, since gene transcription occurs in the same direction for both genes, there is less torsional stress between the two genes and rotation of the downstream gene (mSpinach) enables relaxation of any torsional buildup for itself. However, MG aptamer expression can be adversely affected if mSpinach is actively transcribed, since the viscous drag of the open complex on the mSpinach coding sequence may inhibit free rotation of DNA. Thus, tandem oriented MG aptamer can theoretically experience buildup of positive supercoiling downstream of its coding sequence (and upstream of the Lac promoter), depending on the occupancy of open complexes on the mSpinach coding sequence.

Expression of convergent oriented genes on linear DNA can result in buildup of local torsional stress in between the two genes, with positive supercoiling downstream of the sense cassette and negative supercoiling downstream the anti-sense cassette (see Figure 5). In theory, the negative supercoiling could facilitate expression of the anti-sense gene, while the positive supercoiling would interfere with expression of sense gene. To test this, we amplified linear DNA fragments of mSpinach and MG aptamer in convergent, divergent, and tandem orientation and gel purified each sample. We also amplified single gene linear DNA controls containing either mSpinach or MG aptamer. After an additional PCR purification step (to wash out any salt content from gel purification), we expressed convergent, divergent, and tandem mSpinach and MG aptamer from equimolar concentrations of linear DNA (see Supplemental Figure S1D-F).

Remarkably, we observed that convergent oriented mSpinach expressed significantly higher than divergent, tandem oriented, or the single gene control linear DNA. The expression of both divergent and tandem oriented mSpinach was comparable to the mSpinach control and to each other, suggesting that in the absence of topological barriers, differences in expression between tandem and divergently oriented mSpinach were significantly attenuated.

In contrast, convergent oriented MG aptamer expressed at levels comparable to the control while divergent oriented MG aptamer was expressed at slightly higher concentrations. Most interesting was the complete shutoff of MG aptamer expression in the tandem orientation. This outcome came as a surprise, since we could think of no other hypothesis involving transcriptional interference that could explain the loss of MG aptamer expression. These results, combined with the strong responses of plasmids to gyrase treatment, further validated supercoiling as the source of compositional context effects.

### Supplemental Note: Fitting Hill Functions for Different Gene Orientations

We modify the standard Hill Equation to include a term for promoter leakiness that is independent from the dynamic range due to inducer concentration. The equation for expression due to a promoter with some leakiness and Hill function-type response to an inducer chemical is given as:

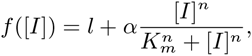

where [I] is inducer concentration, *l* is leaky expression, *α* is the amplitude of expression due to inducer, *n* is the apparent cooperativity of the response to inducer, and *K*_*m*_ is the concentration at which induction is half maximal. Thus, the maximum total expression upon full induction is given by:

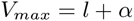

All four parameters were fit using RFP/CFP expression data shown in Figure ??. Both RFP/CFP induction functions were fit for the case in which the other gene is fully induced using the Matlab function `nlinfit`. RFP was fit to the data that varies aTc (1.56 ng/mL to 200 ng/mL) while keeping IPTG at 1000 nM (left column of Figure 4). Similarly, CFP was fit to the data that varies IPTG (7.85 nM to 1000 nM) while keeping aTc at 200 ng/mL (top row of Figure 4). Fits along with experimental data points were plotted for all three orientations for both RFP and CFP (Figure S2A-B).

### Supplemental Note Comparing Toggle Switch Performance of the Original (Divergent) Gardner-Collins Toggle and Convergent Gardner-Collins Toggle

While our experimental results with *de novo* toggle switches showed compositional context can reinforce the toggle’s feedback architecture (Figure 7), we also wanted to compare performance of the convergent toggle with the canonical toggle switch developed by Gardner et al. (2000). To this end, we modified the original pIKE107 toggle switch, which was assembled with divergent oriented LacI and TetR-GFP genes, to include an RFP coding sequence downstream of LacI. All RBSs and promoter sequences were left as originally cloned. Next, we used restriction digest DNA assembly to convert the original toggle switch into a convergent toggle switch. All cloning was done using the pIKE107 plasmid backbone. Plasmids were grown up in cloning strains, sequence verified, and transformed into both MG1655ΔLacI and MG1655 *E. coli.*

Our first experimental test was to confirm that the improved thresholding properties in the convergent toggle were preserved, independent of plasmid backbone identity. We immediately discovered in preliminary experiments that the RBS for TetR-GFP was significantly stronger than the RBS for LacI-RFP, so to draw a fair comparison with our results in Figure 7, we attenuated the concentrations of IPTG by an order of magnitude. We found that convergent oriented Gardner-Collins toggle exhibited strong XOR logic in the high IPTG-high aTC regime and low IPTG-low aTc regime (Supplemental Figure S4A), consistent with our results in Figure 7. In contrast, the divergent Gardner-Collins toggle did not exhibit strong XOR logic in the high IPTG-high aTc regime and the distinction between high GFP-low RFP and high RFP-low GFP states was generally not as clear (see Supplemental FigureS4B).

Our second experimental test was a stability test of the memory properties of the toggle. We found that both in the MG1655ΔLacI *E. coli* and MG1655 *E. coli* strain there were significant differences between the convergent and divergent toggle. In particular, we observed cells transformed with the divergent toggle tended to drift from its high-GFP state into a lower GFP state over time, while the convergent toggle tended to maintain a high-GFP state throughout the course of the entire 53 hour experiment (48 hours post-latching and 5 hours of latching.)

These results can be explained, once again, using our model of supercoiling and its role in strengthening negative feedback in the toggle switch. In the divergent orientation, the supercoiling propagated by proximal promoters generally results in decreased expression of the repressors. This results in weaker repression, since both promoters are affected by the presence of supercoilng (reference the results in Figures 2, 3, 5). This leads to overall reduction in reporter signal, but also leaky repression. As time transpires post-induction (growing in media without inducer), this allows the population of cells to drift from the high-GFP state.

In the convergent orientation, once TetR-GFP is expressed in the high state, it continuously dominates by propagating supercoils into the LacI-RFP transcript region. These supercoils may impose a higher activation threshold for the LacI-RFP state, thus keeping it effectively off throughout the course of the experiment. Remarkably, we observed that even in the presence of constitutively expressed genomic LacI repressor in MG1655 *E. coli*, the convergent toggle did not drift significantly from its initial state. This result can be interpreted as enhanced disturbance rejection capabilities of the convergent toggle; it requires a significant amount of LacI to flip the toggle switch to a high LacI-RFP state. Small amounts of LacI are not sufficient to overcome the combined repression barrier imposed by supercoiling and TetR repression. In this way, the convergent orientation of the toggle reinforces the feedback architecture of the toggle switch, resulting in improved memory, disturbance rejection, and better thresholding performance.

### Deriving Supercoiling Dynamics in a ODE Model of mSpinach and MG Aptamer Expression

In this subsection we explore a detailed model for describing the interplay of supercoiling and gene expression. The motivation to do this arises from 1) experimental results which strongly suggest that supercoiling and not transcriptional interference is the primary cause of differences observed in mSpinach, CFP, RFP, and MG aptamer expression across different gene orientations and 2) the need for a mathematical modeling framework that describes how the temporal dynamics of gene expression vary as a function of supercoiling state and neighboring gene activity.

We consider three structural phenomena that arise in supercoiled DNA: positively supercoiled DNA, negatively supercoiled DNA, and R-loop formation (Drolet, 2006) of the RNAP-DNA elongation complex in negatively supercoiled DNA. We begin with the basic premises of the twin-domain supercoiling models (Liu & Wang, 1987), namely that when a gene is transcribed, negative supercoiling is introduced upstream of the open complex and positive supercoiling is introduced downstream of the open complex. We introduce several concepts from the supercoiling literature (Drolet, 2006; Korbel et al., 2004; Liu & Wang, 1987; Opel & Hatfield, 2001; Rahmouni & Wells, 1992).

**Definition 1** *We define the constant h*_0_ = 10.5 *to be the number of DNA base pairs involved in a single turn of a B-form DNA molecule in its natural state.*

**Definition 2** *We define the linking number α*_*LN*_ *of a region of DNA to be the number of supercoiling turns in that region.*

**Definition 3** *We define the supercoiling density α*_*X*_ *of a region of DNA X of N base pairs length as σ* = *α*_*LN*_/*N*.

Thus, we will assume that the plasmid DNA in our experiments is in its natural B-form configuration. Of course, by simply defining *h*_0_ = 11 or *h*_0_ = 12, it is possible extend our results to consider DNA in its A and Z form respectively.

It is important to note the notions of positive and negative supercoiling correspond to the notions of left-handed twist and right-handed twist, respectively, and are well defined as long as the direction along which gene expression occurs is specified and fixed. For example, a gene expressing in the sense direction (as considered in the model by Wang and Liu (Liu & Wang, 1987) and the recent analysis of Chong and Xie (Chong et al., 2014)) creates *right-handed twist* or negative supercoiling, conformal with the natural twist or direction of turn in a DNA double helix, downstream of the transcription bubble and *left-handed twist*, or positive supercoiling, upstream of the transcription bubble. Thus, the convention that negative-supercoiling builds upstream of gene expression and positive supercoiling downstream, is sensible only when considering ’sense’ transcription.

When a gene expresses in the anti-sense direction, then using the reference frame defined by sense transcription and the right-handed twist of DNA, we note that unless we rotate the axis of the reference frame 180 degrees, the buildup of supercoiling downstream of anti-sense transcription is still *right-handed* (i.e. negative) and the buildup of supercoiling upstream of anti-sense transcription is still *left-handed* (i.e. positive) (see Figure 5 for a visual example).

A simple way to prove this is to construct a physical model of a supercoiled double-helix. Take two ropes, twisted into a double helix with right-handed twist. Note that defining the twist of the double helix as right-handed inherently imposes directionality in your rope (e.g. left to right or bottom to top, your thumb pointing in the direction of right or top). Tie both ends to a topological barrier, e.g. by connecting them to form a loop (like a plasmid) or fused to two separate posts, so that the twist internal to the double helix cannot dissipate past these barriers. Simulate a transcription bubble by pulling the two ropes apart and notice that preceding the bubble (opposing the direction that your thumb pointed) you will see the generation of additional right-handed twist and succeeding the bubble you generate left-handed twist (conformal with the direction of your thumb). Notice that the bubble could have been formed by unwinding the double helix left to right (sense transcription) or right to left (anti-sense transcription). However, it does not matter what direction we unwound the DNA to form the bubble; the end result is the same — negative supercoiling or right-handed twist preceding the bubble and positive supercoiling (or left-handed twist) succeeding the bubble. Thus, the original twist of the DNA, not the direction of bubble propagation, defines what type of supercoiling builds up preceding and succeeding a transcription bubble.

It is important to clarify that we are not declaring the default supercoiling state of of DNA *in vivo* as generally negatively supercoiled. Rather, we are referencing the classical convention that states that the double helix inherently has right-handed curl or twist (Drolet, 2006). Moreover, we make no assertions about the exact amount of additional negative or positive supercoiling introduced surrounding a sense transcription bubble or an anti-sense transcription bubble. Various aspects of the nature of supercoiling build-up and propagation have yet to be characterized fully via experiments, such as the rate of propagation of supercoils, the spatial distribution of supercoils succeeding or preceding a transcription bubble, and how DNA promoter and transcript sequence pertain to the rate at which supercoils are introduced. While our model is thus constructed with the capacity for quantitative prediction, until it is supported by robust estimates of physiological parameters, it is meant provide a mechanistic hypothesis for explaining the effects observed in our *in vivo* and *in vitro* experiments as opposed to exact predictions.

When two genes are present, e.g. in the convergent orientation, the intergenic region between the two genes becomes exposed to both left-handed (positive) and right-handed (negative) twist. It is important to note that left-handed (positive) and right-handed twist (negative) do not simply cancel out — the arbitrary nomenclature of positive or negative twist does not confer the same algebraic consequences of adding positive and negative numbers. Rather, when the a right-handed DNA double helix experiences torsional stress from simultaneously introducing both left-handed and right-handed twist from two opposing point sources (e.g. transcription bubbles in convergent orientation), the two twists define opposing forces that meet each other at some kink point between the two point sources. The outcome is not annihilation of positive and negative supercoiling but rather the transition of the kink along the longitudinal axis of DNA until an *equilibrium* is achieved, i.e. the forces driving left-handed twist through the kink are equally balanced by forces driving right-handed twist through the kink. At equilibrium the *net* force is zero but this does not implicate in any way that the presence of right-handed or left-handed coils have been annihilated. With each transcriptional event or binding of a gyrase or topoisomerase to modulate the surrounding DNA’s supercoiling state, the equilibrium position of the kink is correspondingly adjusted.

With these observations in order, we now consider the scenario when two non-overlapping genes are adjacent to each other in varying orientations. For the purposes of our model, three regions of DNA for each gene will be of interest, the promoter of a transcriptional unit, the coding sequence of a transcriptional unit, and the intergenic spacing region between adjacent genes in our constructs. Supercoiling has been experimentally demonstrated to affect both the processes of transcription initiation and transcription elongation. Thus, we make a point to distinguish and keep track of the supercoiling states of both the promoter and coding transcript. For simplicity of exposition, we do not explicitly model the supercoiling density of the intergenic spacing region, however, our models will implicitly assume that the spacing region is able to absorb supercoils propagated from upstream or downstream transcription events up to the kink (if present).

For notation, when modeling the RNA aptamer plasmids, we will use *TL*_*X*_ where *X* = *G* or *S* to denote the length of the MG and mSpinach RNA aptamer transcript respectively, *EC*_*X*_ to denote the elongation complex formed while transcribing gene *X*, *R* to denote RNA polymerase, *PL*_*X*_ to denote the length of the *p*_Lac_ and *p*_Tet_ promoters, and *N*_*S*_ the length of the intergenic spacing region of noncoding DNA between genes. Similarly, we will use the subscript *X* = *RF* or *CF* to indicate the parameter of interest pertains to the coding sequence for RFP or CFP respectively.

#### Convergent Orientation Model

In the convergent orientation, promoters face each other and as both genes express, positive supercoiling propagates from the sense transcription bubble into the intergenic spacing region and negative supercoiling propagates from the anti-sense transcription bubble (see Figure 4) to form a kink in between the two genes. Where standard transcription translation models of gene expression assume constant rates of transcription initiation and transcription elongation, we now make explicit the dependencies of these rates on supercoiling state. The chemical reaction network for this orientation is given as:

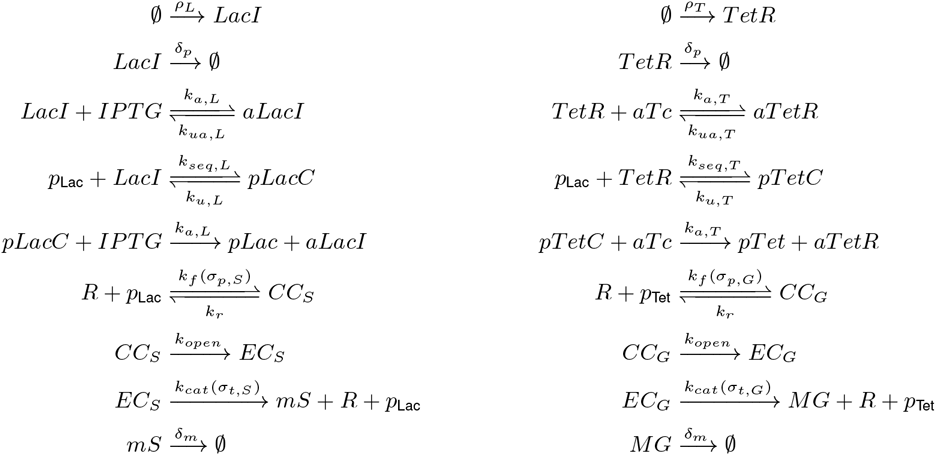

We note here, that for simplicity, we model LacI and TetR as naturally occurring in their tetrameric and dimeric forms. The complex aLacI and aTetR denote the inducer-bound forms of LacI and TetR that are unable to bind to their target promoter. When LacI and TetR bind their respective promoter, we denote them with pLacC and pTetC to indicate the promoter is sequestered from transcriptional processes.

We now derive an expression for *σ*_*p*,*S*_(*t*) by first considering the effects of transcription on the supercoiling density of the transcript. Consider the collection of plasmids present in the cell or volume of cell-free extract. Consider a small time interval [*t*,*t* + *ϵ*] for small *ϵ* > 0. Suppose that Δ_*LN*,*t*_ turns are introduced with the production of each mSpinach transcript and that *x*Δ_*LN*,*t*_ number of turns are introduced into the transcript as *x* mSpinach molecules are produced. Simultaneously, we suppose that if *y* additional open complexes have been formed, then correspondingly *y*Δ_*LN*,*t*_ turns have been removed from the transcript region (in order to facilitate open complex formation). Also, although the promoter region is short, each time a transcription initiation event occurs, the promoter region is unwound and propagates supercoiling. However, many transcription initiation events stall or reversibly dissociate. We suppose such events do not introduce significant amounts of supercoiling — rather, only when an elongation complex is formed do we suppose that the promoter region has been unwound and introduced supercoils in the proximal regions. Thus, we suppose that if there are *y* new elongation complexes, then there are *y*Δ_*LN*,*p*_ turns. Moreover, once the elongation complex departs, it is not necessarily true the promoter will resume its normal B-form DNA state. However, we suppose that the reaction event of a new initiation complex finally forming is indicative of a supercoiling state being removed. In this way, turns are gained and lost by the incoming and outgoing of holoenzyme complexes on the promoter and transcript regions. We assume that any transcriptional pausing, abortive initiation, and aborted elongation events are effectively modeled by their respective transcriptional parameters. The dynamics of *σ*_*t*,*S*_ can then be expressed as:

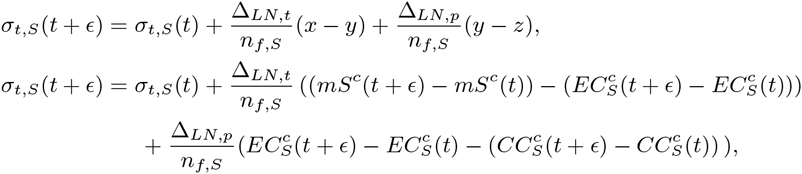

where *σ*_*t*,*S*_(*t*) denotes the supercoiling density at time *t*, *mS*^*c*^(*t*) denotes the integer molecular *count* of total mSpinach molecules produced by time *t*, Δ_*LN*_ denotes the change in the linking number of the mSpinach coding region per mSpinach transcript expressed, and *n*_*f,S*_ is the combined length of free mSpinach transcript and spacer that is able to absorb the residual twist introduced by transcription. The amount of free spacer DNA between negatively supercoiled and positively supercoiled DNA available to absorb additional supercoiling depends on the dynamic equilibrium between negative twist and positive twist from mSpinach and MG aptamer transcription, respectively. We suppose that the length

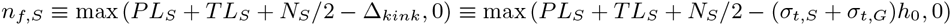

and

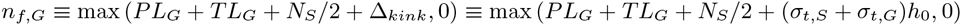

When *σ*_*t*,*S*_ = *σ*_*t*,*G*_ note that the point of transition between negative and positive supercoiling is exactly centered. When *σ*_*t*,*S*_ < *σ*_*t*,*G*_, i.e. the force from negative twist exceeds the force of positive twist, the kink is forced in the direction of the MG aptamer coding transcript and *n*_*f*,*S*_ *>TL*_*S*_ + *N*_*S*_/2. Conversely, if mSpinach transcription does not produce additional negative supercoils to counteract the positive twist from MG aptamer expression, then *n*_*f*,*S*_ *< TL*_*S*_ + *N*_*S*_/2.

The above equation states that the supercoiling density at time *t* + *ϵ* is the supercoiling density at time t with an additive perturbation term, corresponding to the change in supercoiling density from transcription of *x* = *mS*^*c*^(*t* + *ϵ*) − *mS*^*c*^(*t*) transcripts. Normalizing by the reaction volume Ω on both sides, dividing by *ϵ*, and taking *ϵ* → 0, we obtain an expression in terms of the derivative of mSpinach *concentration*:

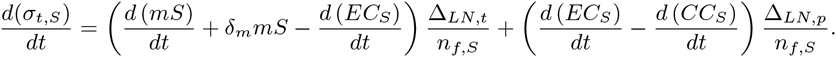

Notice that the quantity *d*(*mS*)/*dt* + *δ*_*m*_*mS* represents the rate at which total mSpinach RNA aptamer is produced in the system, since it is the state dynamics of mSpinach without mRNA degradation. However, the supercoiling state of DNA is continuously regulated by gyrase, an enzyme that relieves positive supercoiling, and topoisomerase, an enzyme that relieves negative supercoiling. We estimate that gyrase relieves positive supercoiling of the transcript region at roughly *γ* = 0.5 turns per second per plasmid, while topoisomerase relieves negative supercoiling of the transcript region at roughly *τ* = 0.25 turns per second per plasmid (Liu & Wang, 1987). Both enzymes act to maintain the natural physiological (negative) supercoiling density of *σ*_0_. We suppose that in the absence of any transcriptional activity, the balance of these rates tends towards gyrase activity and a steady state of σo. For simplicity we suppose that gyrase and topoisomerase binding does not interfere with the transcriptional binding dynamics of polymerase. We incorporate these maintenance dynamics as follows:

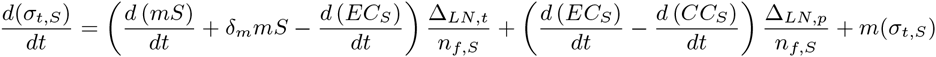

where

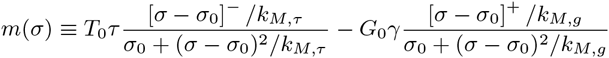

where *v* is the total length of DNA, *x* ≡ [*x*]^−^ + [*x*]^+^ denotes an additive decomposition of x into its strictly negative and nonnegative parts, and *T*_0_ and *G*_0_ are the topoisomerase and gyrase concentrations present *in vivo* or *in vitro* cell-free extract.

Next, to obtain an expression for Δ_*LN*_ *<* 0, i.e. the number of negative supercoiling turns introduced by expression of one mSpinach transcript, we argue as follows. As the open complex proceeds along the anti-sense DNA template of mSpinach, it unwinds and displaces the supercoiling of a 17 base pair region (Liu & Wang, 1987), corresponding to the DNA footprint of a transcription bubble (i.e. DNA-RNAP open complex). The transcription bubble requires an uncoiled region of DNA to transcribe. Thus, an additional 17/*h*_*o*_ turns are introduced into the upstream and downstream regions. We suppose that half of these turns are introduced as negative supercoiling and the other half as positive. Thus, in the wake of the transcription bubble passing through the entire transcript, there are

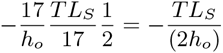

negative supercoiling turns introduced into intergenic spacer downstream. When transcription termination occurs, the bubble is no longer held open by the open complex and the negative supercoils travel back into the unwound DNA of the mSpinach transcript and spacer, while the positive supercoils dissipate upstream of the promoter. Similarly, as the promoter expresses it also introduces negative supercoils downstream into the transcript region. The expression for *σ*_*t*,*S*_(*t*) then simplifies to

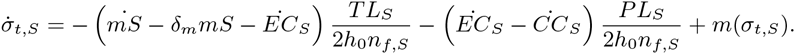

Here we use 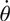 notation to denote the derivative of *θ*. Following similar arguments, we can write the dynamics of *σ*_*p*,*S*_(*t*) as

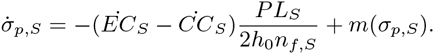

Similarly, the supercoiling density dynamics for the MG RNA aptamer gene are given as:

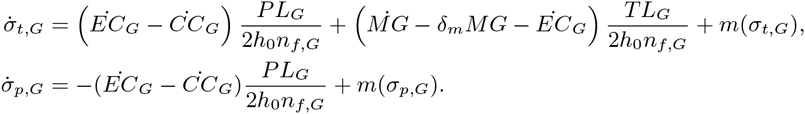

Notice the change in sign in the MG RNA aptamer dynamics. In this way, the directionality of sense transcription, relative to the right-handed twist of DNA, is encoded. If MG RNA aptamer was expressed in the anti-sense direction (which is the case in our divergently orientated construct), then the supercoiling introduced would be negative.

An important question is how transcription initiation rate *k*_*f*_(·) and elongation rate *k*_*cat*_ (·) depends on supercoiling density. In (Meyer et al, 2014) it was argued that the reaction rate of transcription initiation could be modeled with a Hill function type curve, based on experimental data characterizing the pelA and pelE promoters (Ouafa et al., 2012). Although these results are specific to the bacterium *Dickeya dadantii*, it has been generally postulated that supercoiling density acts as a form of global gene regulation (Drolet, 2006; Rahmouni & Wells, 1992) both in prokaryotic and eukaryotic organisms. A study of the ilvY and ilvC promoters (Rhee et al., 1999) in *E. coli* suggest that promoter activity is optimal around around a certain value of *σ*\* and that activity tapers as σ diverges towards positive or negative infinity. Balke and Gralla (Balke & Gralla, 1987) argued that *global* supercoiling state forms the basis of a feedback loop for a system of genes in an organism, in response to environmental cues regarding metabolite and resource availability.

Broadly speaking, it is difficult to draw general conclusions regarding the relation of supercoiling state and promoter activity — all experimental measurements in the studies described above were of the *global* supercoiling density. In these studies, the common approach was to treat a purified plasmid with topoisomerase to introduce additional twist. Whether this twist was introduced uniformly across the plasmid or non-uniformly is unclear. However, we can suppose that when a topoisomerase was used to treat plasmid, it introduced a monotone amount of twist (gyrase introduced *only* negative coils and Topo I introduced *only* positive coils). We thus proceed supposing that incubation and treatment with a topoisomerase had a monotonic effect on promoter supercoiling state and that the overall *qualitative* trends observed regarding the ilvY, ilvC, and pelE, and pelA promoters can be used to inform the *qualitative* or phenomenological model of how *local* supercoiling density and promoter activity are related. Drawing from physical intuition, we argue that a promoter cannot initiate transcription if it is excessively wound with positive or negative twist. We suppose that transcription initiation is thus optimal at a particular value of local supercoiling density *σ*\*. Moreover, we suppose that for a given promoter of length *PL*_*X*_, *X* = *S,G, RF*, or *CF* the optimal *local* supercoiling density optimum roughly is related to the optimal *global* supercoiling density *σ*_0_ via the following approximation:

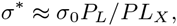

where *P*_*L*_ is the length of the plasmid. We model the rate of transcription initiation as a second-order symmetric Hill function, with an optimum centered around *σ*\*.

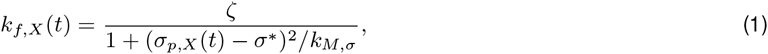

where *X* = *G* or *S* for MG and mSpinach transcription respectively and ζ is the optimal putative forward reaction rate of transcription initiation assuming the supercoiling state *σ*_*p,X*_ is optimal for transcription initiation. Similarly, we suppose in the case of transcriptional elongation that the optimum *σ*\* = *σ*_0_*P*_*L*_/*TL*_*X*_ and the elongation rate is defined by the functions

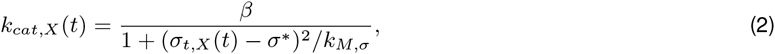

where *X* = *G* or *S* for MG and mSpinach respectively and *β* is the putative transcription elongation rate when the supercoiling state *σ*_*t,X*_ is optimal for transcription. Finally, we note the following conservation laws hold since DNA and RNAP concentration are constant in our *in vitro system*

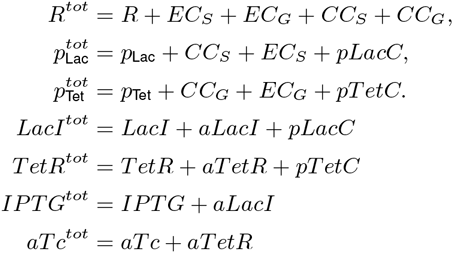

Using these laws, we can write a reduced order dynamical system model for the *convergent* biocircuit:

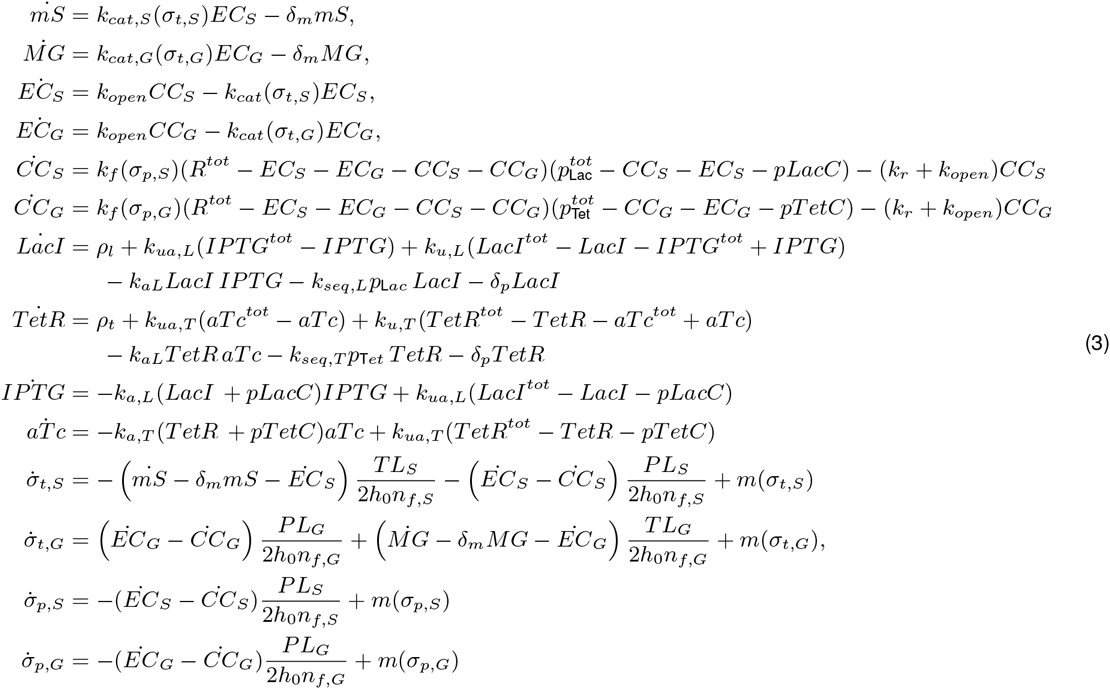

In simulating the supercoiling dynamics we noticed that the magnitude of our *local* supercoiling states settle around steady-state values much higher than the traditionally accepted range of global supercoiling density. In practice, experiments have determined that DNA is negatively supercoiled with a *global* supercoiling density of −0.065 and can drop to as low as −0.1. This parameter does not reflect the *local* supercoiling density of the regions of interest in our model, namely the supercoiling density of the transcript and the promoter.

For example, a small region of DNA can maintain a positively coiled plectonome while the rest of the DNA is relatively relaxed. The global supercoiling density will reflect the overall twist, as opposed to the high density in either of the two regions. Wang and Liu estimated that expression of a single transcript can introduce supercoils into DNA at a rate of 4 supercoils per second per transcript. Assuming gyrase introduces γ = 1 negative supercoils per second and τ = .5 positive supercoils per second on a given plasmid, if half the supercoils introduced propagate upstream and the over half downstream, over the course of just five minutes (Liu & Wang, 1987) the region downstream (such as the 150 bp spacer sequence in our plasmids) of the transcript could achieve a *local* supercoiling density of *σ* = 2.0. A measurement of the *global* supercoiling state of the plasmid, say 3.5 kbp in length, would yield a *global* estimate of only *σ* = 0.08! Therefore, it is important to note the distinction between *local* and *global* supercoiling density; the *local* supercoiling density of a region of DNA can reach much higher magnitudes despite a relatively low (and conventionally acceptable) *global* supercoiling density.

### Divergent Orientation Model

In our divergently oriented plasmid, the Tet promoter and Lac promoter express in opposing directions, but the transcription bubbles diverge or move away from each other. Thus, the only torsional stress introduced comes from backward propagation of coils from unwinding the regions of DNA encoding promoters into the intergenic spacing region between the two genes. The Tet promoter back propagates positive supercoils into the intergenic spacing region while the Lac promoter back propagates negative supercoils. The position of dynamic equilibrium between the positively supercoiled region upstream of the Tet promoter and the negatively supercoiled region upstream of the Lac promoter is determined by the balance of forces arising from positive and negative twist (diametrically opposing each other) in the promoter supercoiling states *σ*_*p,S*_ and *σ*_*p,G*_. If *σ*_*p,S*_ is much larger than *σ*_*p,G*_ then the equilibrium shifts in favor of the Lac promoter and the positive coils are pushed closer to the actual Tet promoter (or vice-versa). We model the amount of spacer available to the promoters as *n*_*f,S*_ and *n*_*f,G*_ where

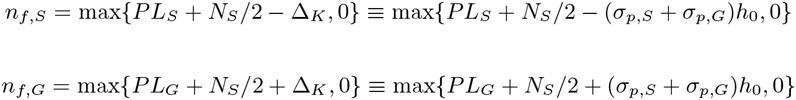

again noting that *n*_*f,S*_ + *n*_*f,G*_ = *PL*_*S*_ + *PL*_*G*_ + *NS* base pairs defines the total length of DNA in which localized supercoiling buildup can propagate. We suppose that all other supercoils arising from transcription elongation are dissipated within regions downstream of the promoters.

The supercoiling dynamics of the divergently oriented construct for mSpinach and MG RNA aptamer are thus given as

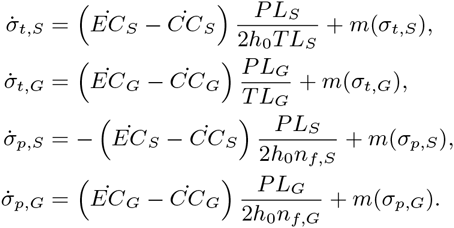

while the rest of the system dynamics are as presented in the convergent model. Any differences in expression are thus a function of the supercoiling dynamics above, the initial conditions of these four states, and their effects on *k*_*f*_, *k*_*cat*_ and the topoisomer maintenance function *m*(·).

### Tandem Orientation Model

In the tandem orientation, negative supercoiling backpropagates from the *p*_Lac_ promoter into the intergenic spacing region between the MG aptamer coding sequence and the mSpinach promoter. The torsional stress introduced by downstream propagation of positive supercoils from MG aptamer elongation and upstream propagation of negative supercoils from mSpinach transcription initiation again defines a dynamic equilibrium that is determined by the balance of *σ*_*t,G*_ and *σ*_*p,S*_. When the Lac promoter for mSpinach is much more active relative to the transcriptional activity of the MG aptamer coding sequence, *σ*_*p,S*_ can dominate *σ*_*t,G*_ such that any residual positive supercoils from MG aptamer transcription are pushed back into the coding sequence for MG aptamer. Excessive negative supercoiling from the mSpinach promoter, likewise, can make their way into the MG aptamer coding sequence. This is especially likely if the transcript region of MG aptamer is short (since it generates less positive supercoils to counteract the twisting force of negative supercoiling from mSpinach expression). The presence of excessive negative supercoiling in a transcript region can result in the formation of a R-loop complex, a hybrid of the RNAP-DNA open complex and the nascent mRNA chain with upstream DNA (Drolet, 2006); this complex stalls the elongation process indefinitely and impedes subsequent transcription events. These effects are accounted for in the function *k*_*cat*_(*σ*), which tapers off towards 0 if *σ* → −∞. The sensitivity of *k*_*cat*_ to -*σ* is determined by the parameter *k*_*M,σ*_.

Alternatively, if MG aptamer expression is high or leaky, it can likewise propagate positive supercoils downstream into the spacer region, which subsequently shutoff promoter activity of mSpinach. The decrease in promoter activity in mSpinach only further enables MG aptamer expression, which leads to MG aptamer dominant expression. This is particularly relevant if the coding sequence for MG aptamer is long, or replaced with a long coding sequence for a protein, e.g. RFP. In such a scenario, the expression of RFP can repress future expression of mSpinach.

It is thus important to consider the dynamic equilibrium of *σ*_*t,G*_ and *σ*_*p,S*_ and how the balance of these forces impact the positioning of positive and negative supercoils in the transcript region of MG aptamer and the Lac promoter. We suppose that the length of DNA available for positive supercoiling buildup (from MG aptamer expression) is given as:

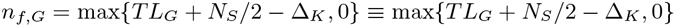

and the length of DNA available for negative supercoiling buildup (from mSpinach expression) is given as:

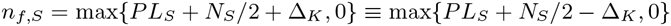

The supercoiling dynamics are thus given by

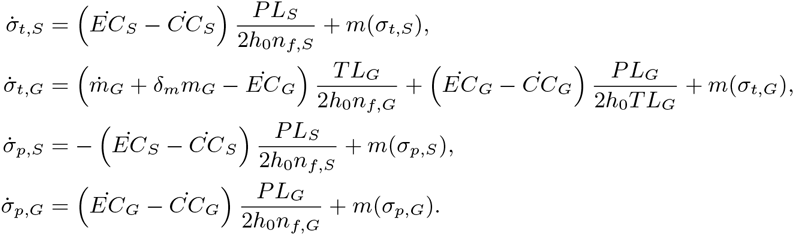

#### RFP CFP reporter models

Modeling the expression of RFP and CFP instead of mSpinach and RFP does not change any of the preceding arguments for deriving dynamics of supercoiling states. The only difference at the transcriptional level is that we use different length parameters, *TL*_*R*_ and *TL*_*C*_ (see Table 1) to define the length of transcript regions and denote the transcriptional products of each transcription elongation reaction as *m*_*C*_ (for the CFP mRNA transcript) and *m*_*R*_ (for the RFP mRNA transcript).

**Table 1:** Parameters used for the deterministic ODE model for convergent, divergent, and tandem oriented reporters

| Parameter Name | Description | Numerical Value | Source |
| --- | --- | --- | --- |
| $h_0$ | Bps per right-handed turn in B-DNA | 10.5 bp/turn | Berg et al. |
| $TL_S$ | mSpinach-tRNA and T500 term. length | 203 bp | Larson et al.; Paige et al. |
| $TL_G$ | MG aptamer and T500 terminator transcript length | 68 bp | Babendure et al. |
| $N_S$ | Intergenic spacer length | 150 bp | NA |
| $PL_S$ | Lac promoter length | 40 bp | Lutz & Bujard |
| $PL_G$ | Tet promoter length | 44 bp | Lutz & Bujard |
| $p_{Len}$ | Length of ColE1-mSpinach-MG reporter plasmid | 2892 bp | NA |
| $K_{M,\sigma}$ | Michaelis constant for supercoiling Hill functions | 50 nM | NA |
| $R^{tot}$ | Total RNAP concentration | 18.931 μM (RNAP) | Bremer & Dennis |
| $Ribo^{tot}$ | Total Ribosome concentration | 11.291 μM (ribosome) | Bremer & Dennis |
| $p_{Lac}^{tot}$ | Total Plasmid Concentration | 11 nM | NA |
| $p_{Tet}^{tot}$ | Total Plasmid Concentration | 11 nM | NA |
| $G_0$ | Gyrase Concentration | 12 nM | Maier et al. |
| $T_0$ | Topoisomerase Concentration | 2 nM | Maier et al. |
| $LacI^{tot}$ | LacI concentration | 10 nM | Kalisky et al. |
| $TetR^{tot}$ | TetR concentration | 0 nM (TX-TL) | Kalisky et al. |
| $IPTG^{tot}$ | IPTG concentration | 0 nM (TX-TL) | NA |
| $aTc^{tot}$ | aTc concentration | 0 nM (TX-TL) | NA |
| $k_{f,max,l}$ | Leaky forward transcription initiation rate | $7 \times 10^{-2}$ nM/s | Siegal-Gaskins et al. |
| $k_r$ | Reverse transcription initiation rate | 550 $s^{-1}$ | Bintu et al. |
| $k_{cat,max}$ | mSpinach-MG averaged transcription rate | $k_{cat,g}/(105.5)$ | NA |
| $k_{cat,g}$ | Per base-pair transcription rate | 85 nt/s | Bremer & Dennis |
| $k_{open}$ | Rate of open complex formation | 0.04/s | Buc & McClure |
| $k_l$ | Fraction of terminator-escaped transcripts | 0.02 | Larson et al. |
| $\rho_l$ | Constitutive production rate of LacI | 0 nM/s (TX-TL) | NA |
| $\rho_t$ | Constitutive production rate of TetR | 0 nM/s (TX-TL) | NA |
| $k_{TL}^0$ | Averaged translation rate of RFP/CFP | $21/(TL_C + TL_R)$ /s | Bremer & Dennis |
| $k_{f,C}$ | Folding rate of CFP | $1/(30 \times 60)$/s | NA |
| $k_{f,R}$ | Folding rate of RFP | $1/(110 \times 60)$/s | Zhang et al. |
| $k_{B16}$ | Relative Ribosomal Affinity | 0.75 | NA |
| $k_{B9l}$ | Relative Ribosomal Affinity | 1.25 | NA |
| $\delta_{m,S}$ | mSpinach degradation rate | $\log(2)/(30 \times 60)$ /s | NA |
| $\delta_{MG}$ | MG degradation rate | $\log(2)/(60 \times 60)$ /s | NA |
| $\tau$ | Negative coils introduced per sec. per Topol | 0.5 /s | Liu & Wang |
| $\gamma$ | Positive coils introduced per sec. per Gyrase | 0.5 /s | Liu & Wang |
| $\sigma_0$ | Natural superhelical density of DNA | -0.065 | Rhee et al. |
| $k_{M,gyr}$ | Hill constant for Gyr. Maintenance Function | 200 nM | NA |
| $k_{a,L}$ | IPTG binding rate to free/DNA bound LacI | $6 \times 10^3$ /s | Kalisky et al. |
| $k_{ua,L}$ | IPTG-LacI disassociation rate | 1 /s | Xie et al. |
| $k_{seq,L}$ | Binding rate of LacI to promoter | 10 /s | Kalisky et al. |
| $k_{u,L}$ | Fall off rate of LacI to DNA | 0.022 /s | Nelson & Sauer |
| $k_{a,T}$ | aTc binding rate to free and DNA bound TetR | $k_{a,L}$ | NA |
| $k_{ua,T}$ | aTc-TetR disassociation rate | $k_{ua,L}$ | NA |
| $k_{seq,T}$ | Binding rate of TetR to promoter | $k_{seq,L}$ | NA |
| $k_{u,T}$ | Fall off rate of TetR to DNA | $k_{u,L}$ | NA |
| $\delta_p$ | Degradation rate for untagged proteins | 0/s (TX-TL) | Shin & Noireaux |

**Table 2:**
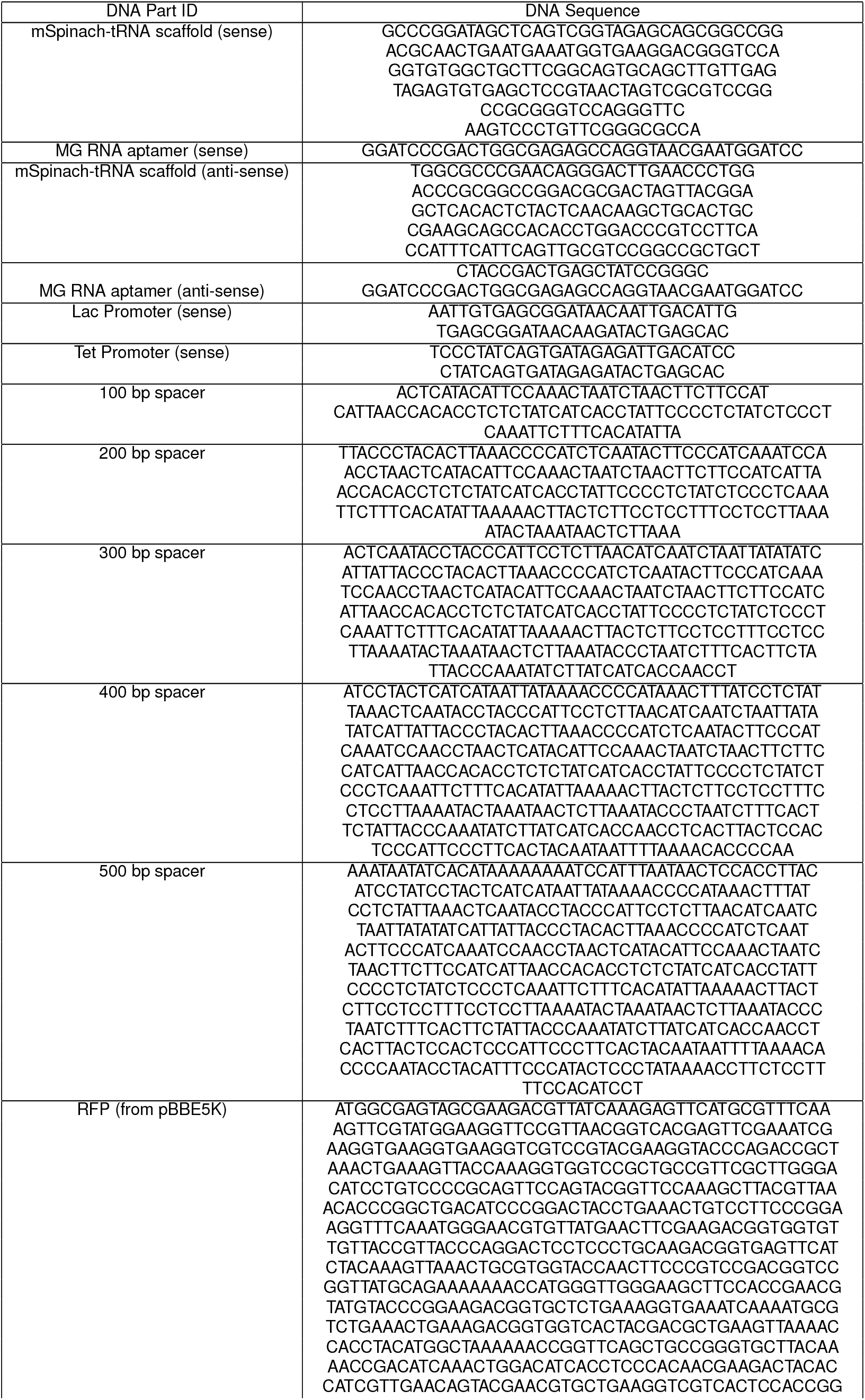
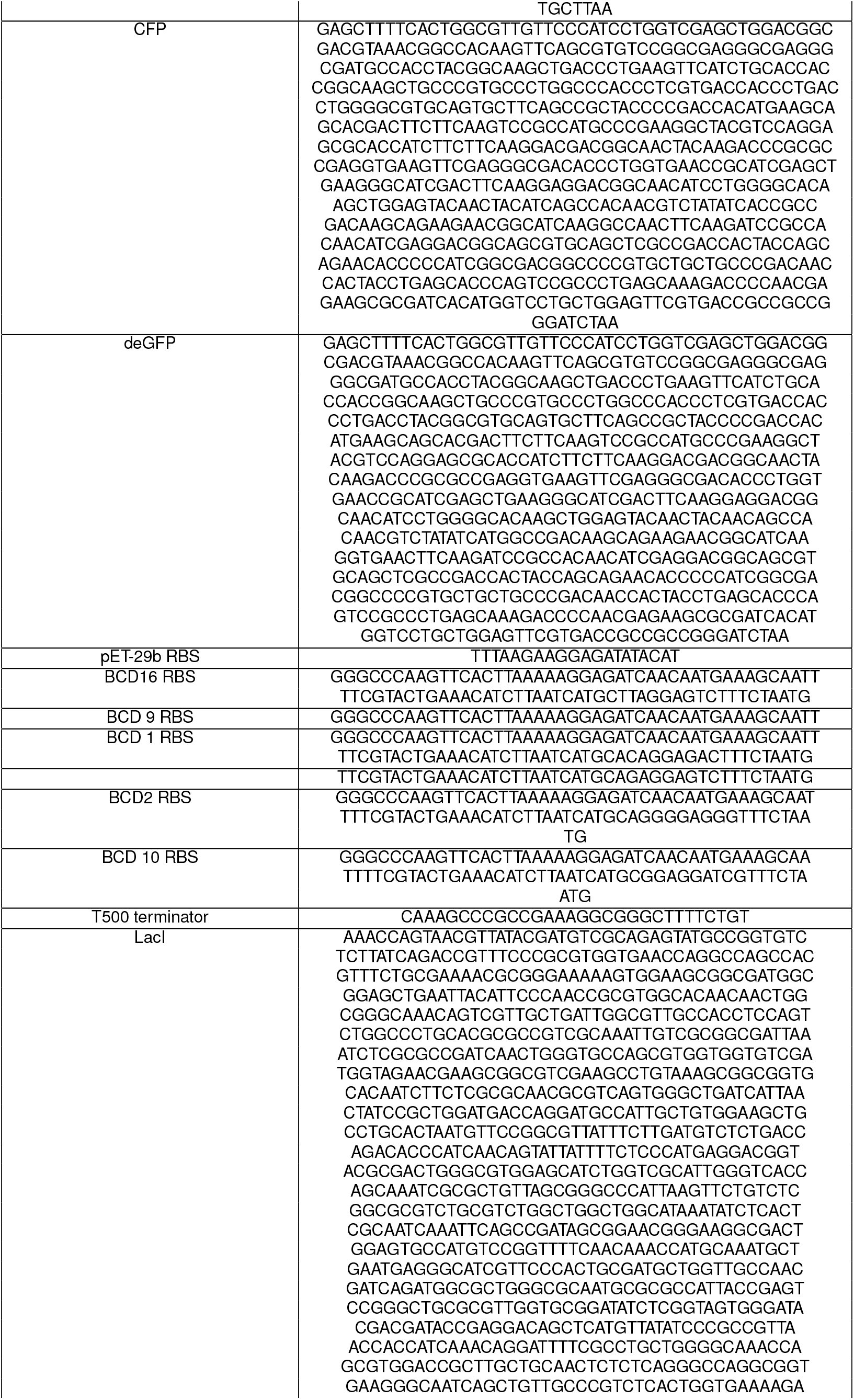
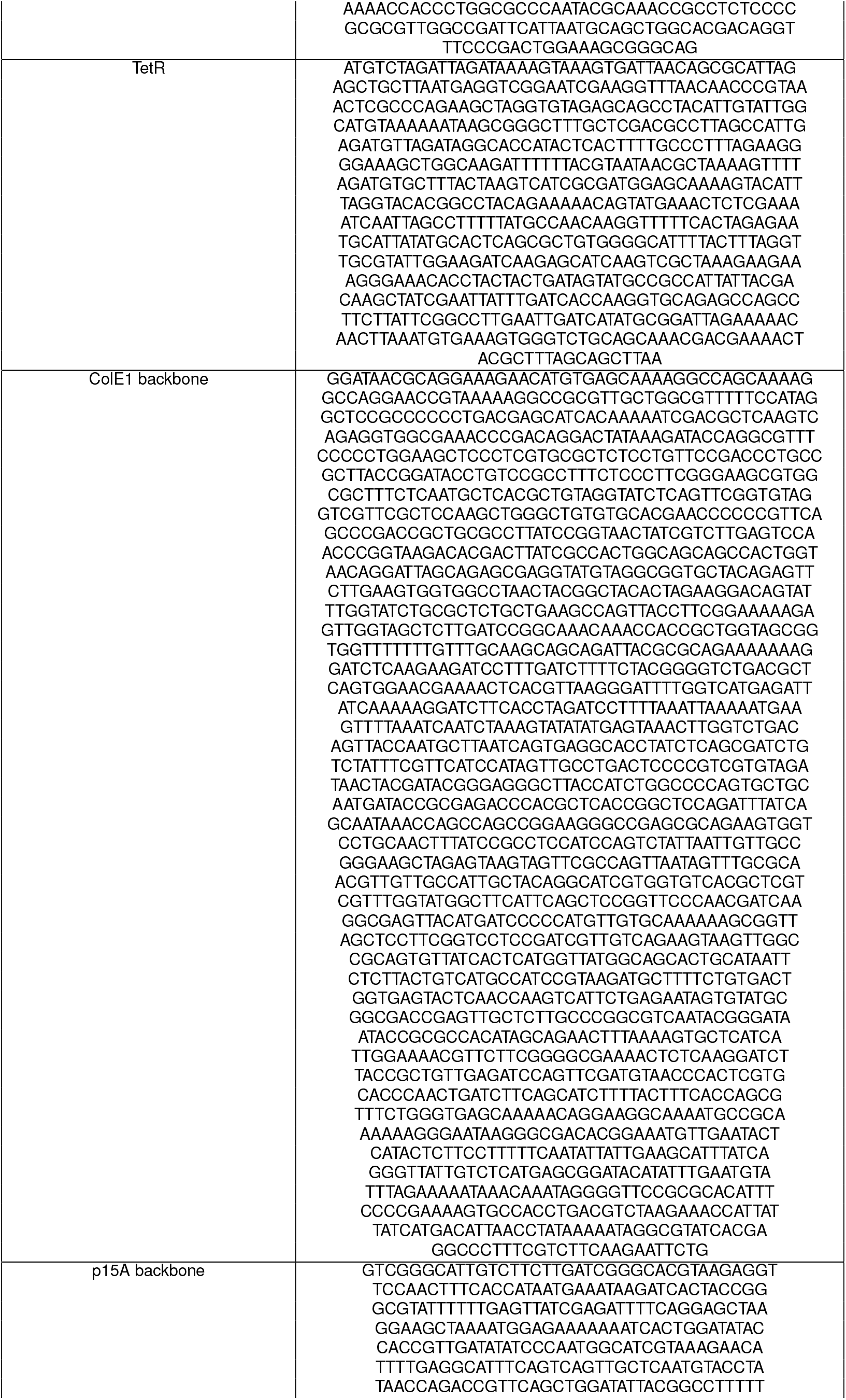
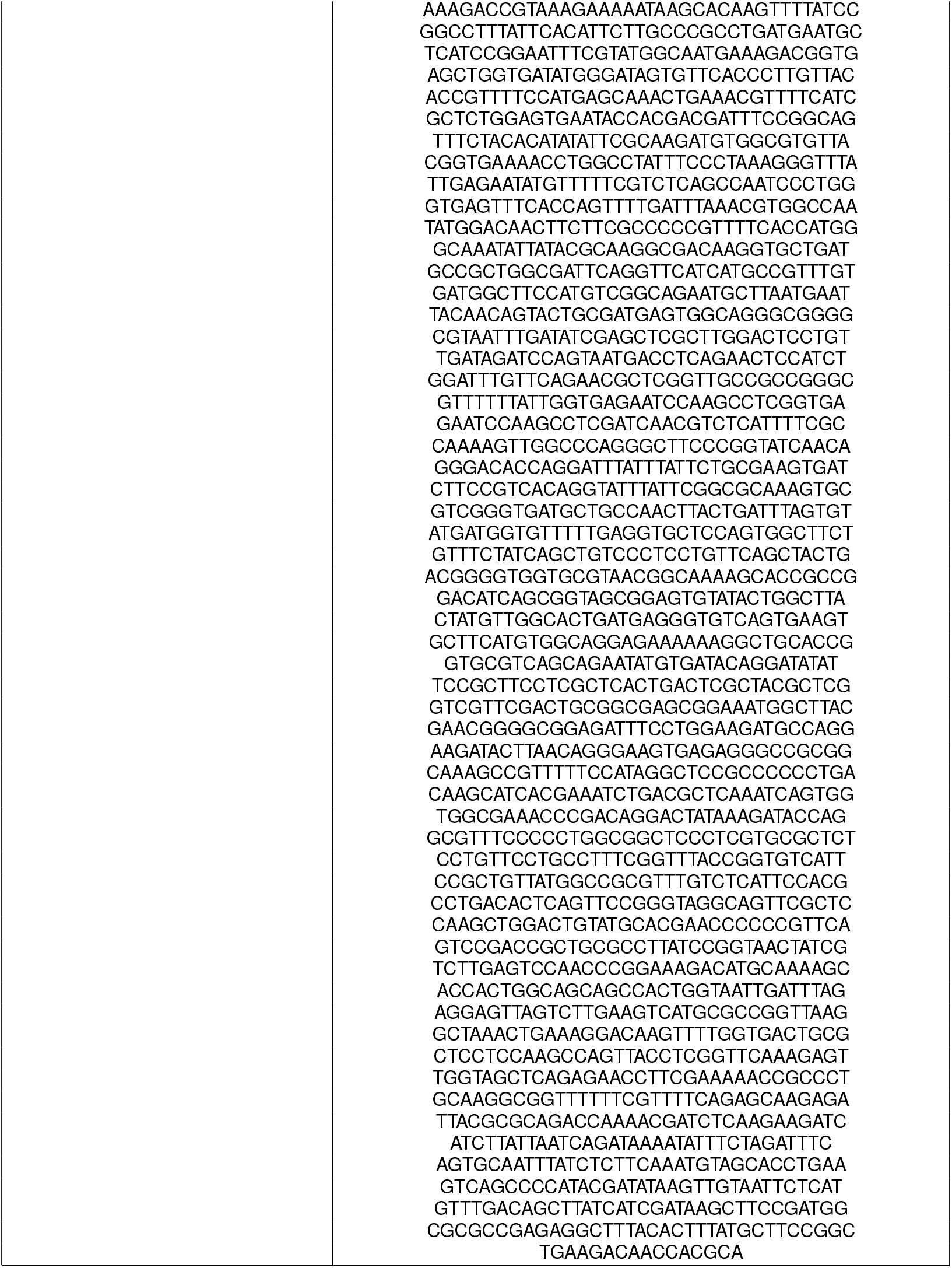
A list of part IDs and DNA sequences used for constructing reporter and toggle-switch plasmids for this paper.

The primary source of genetic context effects in our model is supercoiling at the DNA level. Therefore, in this work we do not consider the effects of secondary structure in mRNA or superhelicity of mRNA-DNA hybrids. Thus, we deliberately model translation reactions simplistically, with the following chemical reactions:

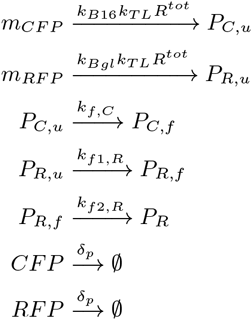

We suppose these translation reactions are the same for convergent, divergent, and tandem oriented genes. Notice the inclusion of maturation reactions for CFP and RFP. We suppose that CFP matures through a one-step process while RFP matures through a two-step process (Zhang et al., 2002). Here we do not necessarily assume that RFP is dimeric, since the variant of dsRed1 that we used in our experiments is actually monomeric. However, we suppose that there is an intermediate stage between unfolded RFP and the final folded RFP We found that including this intermediate stage recapitulated the significant delay observed in RFP expression in the cell-free expression system, that was not seen in CFP

Moreover, the cell-free expression system is typically run in a bulk reaction setting, as a closed biochemical reaction system with a finite and limited amount of ATP, NTPs, and energy molecules to carry out transcription and translation. It has been observed empirically and shown through experiments that as ADP levels build up relative to ATP, enzymatic reactions become increasingly unfavorable (otherwise fluorescent reporters not subject to degradation would express in unbounded and increasing concentrations). Throughout our experiments, we observed these effects of resource depletion, beginning at *t*_0_ ≈ 2 hours onwards. To be consistent with the modeling approaches of Tuza and Singhal, (Siegal-Gaskins et al., 2014; Tuza et al., 2013), we suppose that the translation rate *k*_*TL*_(*t*) decays with time as a first order process, beginning at time *t*_0_, and with decay parameter *α*_*d*_ = log(2)/(480)

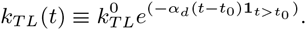

where 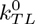 is the nominal translation rate assuming an open system with limitless ATP and energy.

The additional reaction dynamics in the state-space model are thus specified as follows:

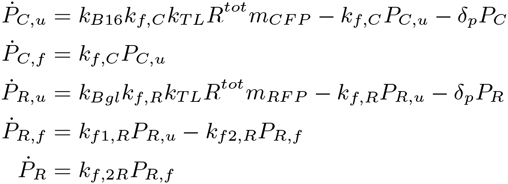

The outcomes of our simulations are plotted in Figure 5 using parameters from Table 1. We see that RFP and CFP expression varies depending on orientation, initial condition of supercoiling states, and that the model is able to recapitulate the trends observed in the data.

It is important to remark that while our model is able to describe the effects observe, it is the gyrase experiments that definitively confirm the validity of supercoiling as a working hypothesis for the physical mechanism driving compositional context effects. Our model serves to validate supercoiling as a hypothesis for compositional context, but not necessarily to prove it.

In conclusion, we have constructed three versions of a simple biocircuit to motivate the need to model compositional context in biocircuit assembly. Our initial data suggests that promoter orientation between pairs of promoters has a salient effect on gene expression. We developed a nonlinear model incorporating various phenomena resulting from compositional context and show it captures the patterns seen in experiments. We emphasize that these results are wholly the consequences of compositional context. There is no designed interaction in the biocircuit, yet different expression biases arise depending on how genes are arranged. Therefore, with any biocircuit comprised of multiple parts, modeling the effects of compositional context should be a chief consideration during the design and prototyping process.

**Figure S1:**
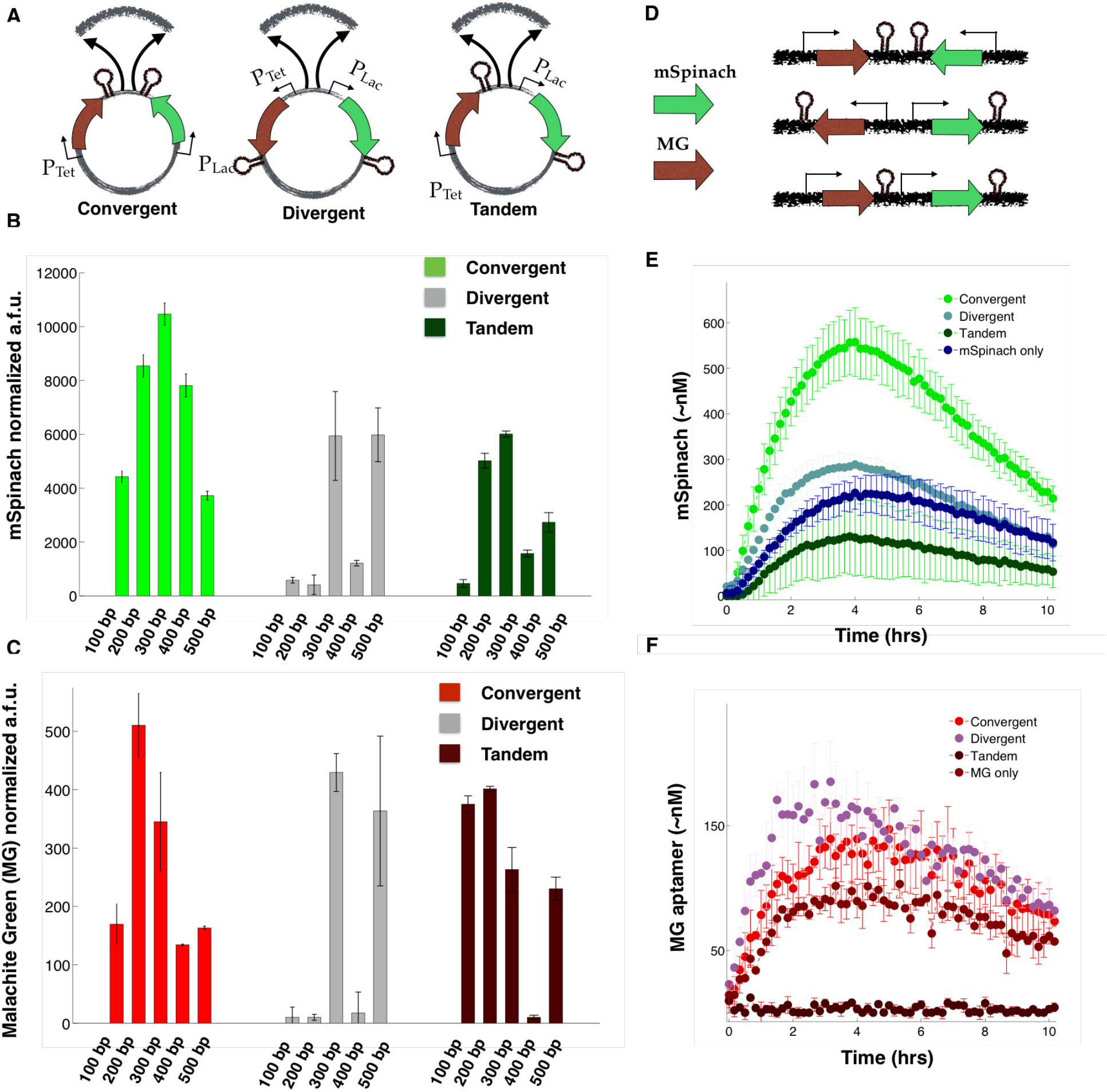
Related to Figure 2. **(A)** A schematic showing the point of insertion of intergenic spacing sequences of length *n* = 100,200,300,400, and 500 bp. **(B)** Steady-state *in vivo* expression of mSpinach from overnight induction in 1 mM IPTG and 200 ng/mL aTc in convergent, divergent, and tandem orientation, varied as a function of spacer length. **(C)** Steady-state expression of MG RNA aptamer from overnight induction in 1 mM IPTG and 200 ng/mL aTc in convergent, divergent, and tandem orientation, varied as a function of spacer length. **(D)** Diagram of linear DNA fragments with mSpinach and MG RNA aptamers in convergent, divergent, and tandem orientation. **(E)** Cell-free *in vitro* expression of equimolar concentrations of linear mSpinach in convergent, divergent, tandem orientation and as a single gene on a linear DNA. **(F)** Cell-free *in vitro* expression of equimolar concentrations of linear MG RNA aptamer in convergent, divergent, tandem orientation, and as a single gene on linear DNA.

**Figure S2:**
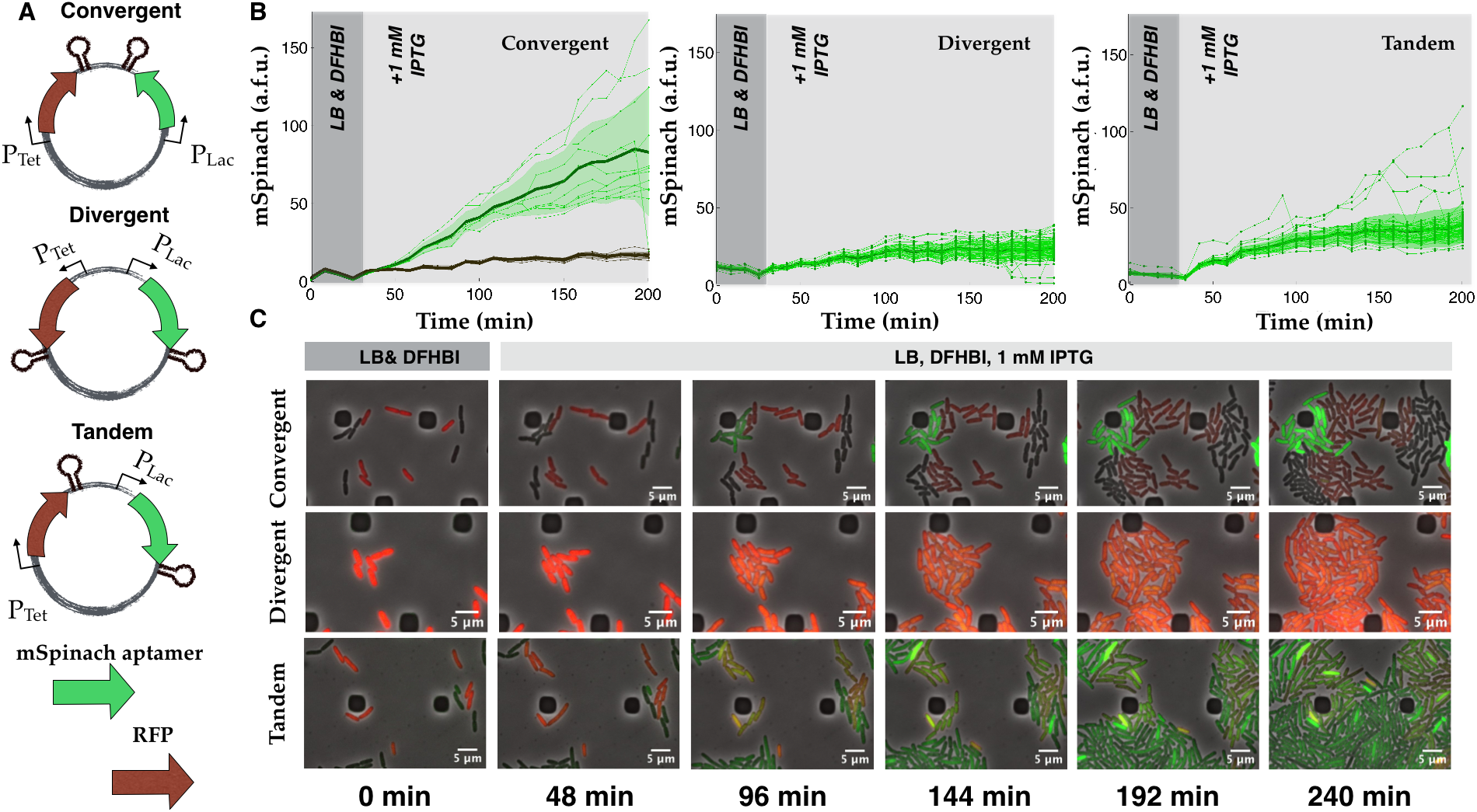
Related to Figures 2 and 3. **(A)** Convergent, divergent, and tandem oriented mSpinach and RFP reporters on ColE1 backbone. **(B)** Time-lapse mSpinach expression curves for individual cell traces in response to 1 mM IPTG induction. Notice that even though mRFP is not induced, its presence significantly affects the magnitude and shape of gene expression. **(C)** Single cell microscopy images of convergent oriented (top) mSpinach and RFP expression, cells responded with a strong bimodal phenotype, (middle) divergent oriented RFP and mSpinach, and (bottom) tandem oriented RFP and mSpinach.

**Figure S3:**
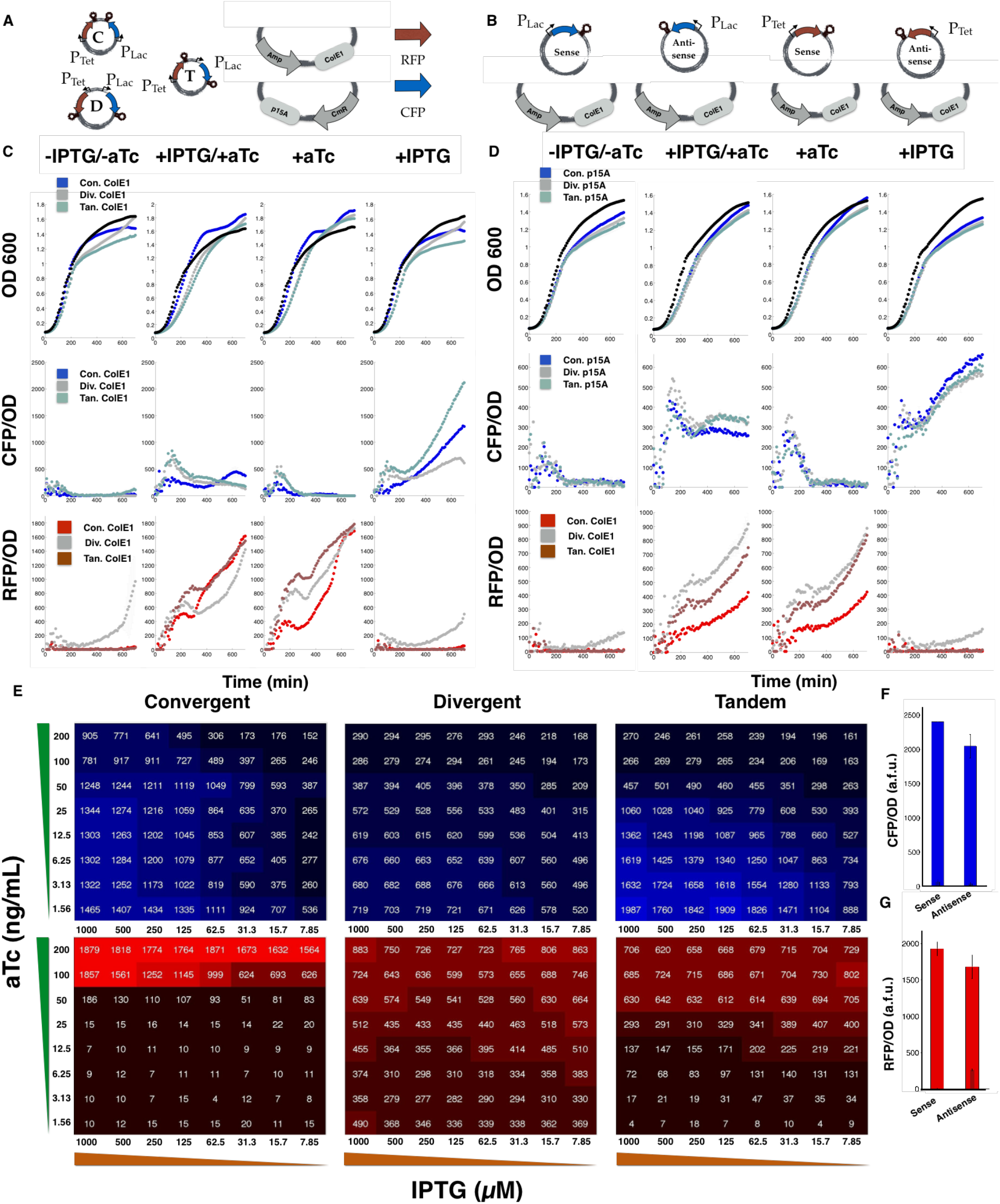
Related to Figures 3 and 4. **(A)** Plasmid layouts for RFP and CFP in convergent, divergent, and tandem orientation, the composition of the plasmid backbone for the ColE1 and p15A backbones used in collecting data for **C-D**. **(B)** A diagram showing the sense and anti-sense CFP and RFP single gene cassette controls, expressed on the ColE1 backbone. **(C-D)** Time lapse *in vivo* plate reader expression of RFP and CFP and growth curves, induced with either 1 mM IPTG, 200 ng/mL aTc, or both, on either ColE1 plasmid or p15A plasmid backbone. **(E)** Quantitative heat-maps of CFP and RFP expression in two variable titration assays of IPTG and aTc for convergent, divergent, and tandem oriented ColE1 plasmids (Figure 4). IPTG is titrated left to right with 2x dilutions starting from 1 mM IPTG (far left) while aTc is titrated top to bottom with 2x dilutions starting from 200 ng/mL. **(F-G)** Expression at *t* = 550 minutes for CFP and RFP in sense and anti-sense orientation.

**Figure S4:**
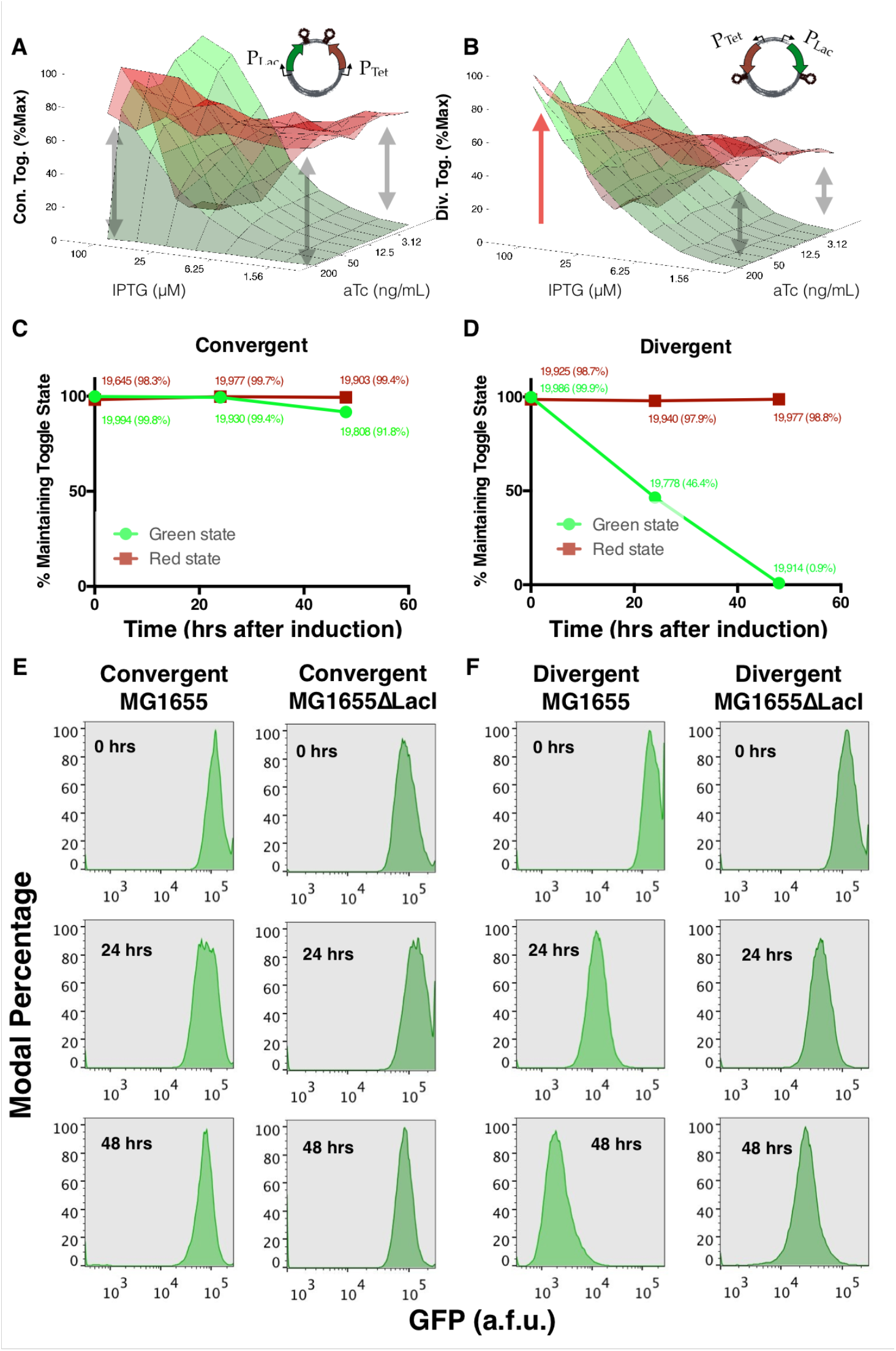
Related to Figure 7. **(A)** Experimental data from a dual reporter expression assay, titrating both IPTG and aTc concentrations to evaluate threshold behavior of the convergent Gardner-Collins toggle switch in MG1655ΔLacI *E. coli.* **(B)** Experimental data from a dual reporter expression assay, titrating both IPTG and aTc concentrations to evaluate threshold behavior of the divergent Gardner-Collins toggle switch in MG1655ΔLacI *E. coli.* **(C-D)** A stability test of the original Gardner-Collins toggle switch and its convergent counterpart in MG1655 *E. coli*. Cells were latched for 24 hours prior to the start of the experiment (*t* = −24 to *t* = 0) and subsequently rediluted in inducer-free media to assess stability of the toggle. The fraction of cells maintaining the original on-state are plotted against time. **(E)** Distributions showing stability of convergent toggle in the high GFP state in cell populations of MG1655 *E. coli* and MG1655ΔLacI *E. coli* plotted at *t* =0,24, and 48 hours. **(F)** Distributions showing stability of divergent toggle in the high GFP state in cell populations of MG1655 *E. coli* and MG1655ΔLacI *E. coli* plotted at *t* = 0, 24, and 48 hours.

## REFERENCES

J. R. Babendure, et al. (2003). ‘Aptamers switch on fluorescence of triphenylmethane dyes’. Journal of the American Chemical Society 125(48): 14716–14717.

V. L. Balke & J. Gralla (1987). ‘Changes in the linking number of supercoiled DNA accompany growth transitions in Escherichia coli.’. Journal of Bacteriology 169(10): 4499–4506.

J. M. Berg, et al. (2002). Biochemistry. Freeman and Company: New York.

L. Bintu, et al. (2005). ‘Transcriptional regulation by the numbers: models’. Current Opinion in Genetics & Development 15(2): 116–124.

H. Bremer & P. P. Dennis (1996). Modulation of chemical composition and other parameters of the cell by growth rate., chap. 97. Springer.

H. Buc & W. R. McClure (1985). ‘Kinetics of open complex formation between Escherichia coli RNA polymerase and the lac UV5 promoter. Evidence for a sequential mechanism involving three steps’. Biochemistry 24(11): 2712–2723.

S. Cardinale & A. Arkin (2012). ‘Contextualizing context for synthetic biology-identifying causes of failure of synthetic biological systems’. Biotechnology Journal.

F. Ceroni, et al. (2015). ‘Quantifying cellular capacity identifies gene expression designs with reduced burden’. Nature methods 12(5): 415–418.

J. Chappell, et al. (2013). ‘Validation of an entirely in vitro approach for rapid prototyping of DNA regulatory elements for synthetic biology’. Nucleic Acids Research 41(5): 3471–3481.

S. Chong, et al. (2014). ‘Mechanism of transcriptional bursting in bacteria’. Cell 158(2): 314–326.

R. S. Cox, et al. (2007). ‘Programming gene expression with combinatorial promoters’. Molecular Systems Biology 3(1): 145.

J. H. Davis, et al. (2011). ‘Design, construction and characterization of a set of insulated bacterial promoters’. Nucleic Acids Research 39(3): 1131–1141.

D. Del Vecchio, et al. (2008). ‘Modular cell biology: retroactivity and insulation’. Molecular Systems Biology 4(1): 161.

M. Drolet (2006). ‘Growth inhibition mediated by excess negative supercoiling: the interplay between transcription elongation, R-loop formation and DNA topology’. Molecular Microbiology 59(3): 723–730.

A. D. Edelstein, et al. (2014). ‘Advanced methods of microscope control using *µ*Manager software’. Journal of Biological Methods 1 (2).

M. B. Elowitz & S. Leibler (2000). ‘A synthetic oscillatory network of transcriptional regulators’. Nature 403(6767): 335–338.

M. B. Elowitz, et al. (2002). ‘Stochastic Gene Expression in a Single Cell’. Science 297(5584): 1183–1186.

C. Engler, et al. (2008). ‘A one pot, one step, precision cloning method with high throughput capability’. PloS ONE 3(11): e3647.

G. S. Filonov, et al. (2015). ‘In-gel imaging of RNA processing using Broccoli reveals optimal aptamer expression strategies’. Chemistry & Biology 22(5): 649–660.

T. S. Gardner, et al. (2000). ‘Construction of a genetic toggle switch in Escherichia coli’. Letters to Nature 403.

D. G. Gibson, et al. (2010). ‘Creation of a bacterial cell controlled by a chemically synthesized genome’. science 329(5987): 52–56.

D. G. Gibson, et al. (2009). ‘Enzymatic assembly of DNA molecules up to several hundred kilobases’. Nature Methods 6(5): 343–345.

A. Gyorgy, et al. (2015). ‘Isocost lines describe the cellular economy of genetic circuits’. Biophysical Journal 109(3): 639–646.

K. Y. Han, et al. (2013). ‘Understanding the Photophysics of the Spinach-DFHBI RNA Aptamer-Fluorogen Complex To Improve Live-Cell RNA Imaging’. Journal of the American Chemical Society 135(50): 19033–19038.

T. Kalisky, et al. (2007). ‘Cost-benefit theory and optimal design of gene regulation functions’. Physical Biology 4(4): 229.

H. Kobayashi, et al. (2004). ‘Programmable cells: interfacing natural and engineered gene networks’. Proceedings of the National Academy of Sciences of the United States of America 101(22): 8414–8419.

J. O. Korbel, et al. (2004). ‘Analysis of genomic context: prediction of functional associations from conserved bidirectionally transcribed gene pairs’. Nature Biotechnology 22(7): 911–917.

S. Kosuri, et al. (2013). ‘Composability of regulatory sequences controlling transcription and translation in Escherichia coli’. Proceedings of the National Academy of Sciences 110(34): 14024–14029.

M. H. Larson, et al. (2008). ‘Applied force reveals mechanistic and energetic details of transcription termination’. Cell 132(6): 971–982.

M. E. Lee, et al. (2015). ‘A highly characterized yeast toolkit for modular, multipart assembly’. ACS Synthetic Biology.

T. S. Lee, et al. (2011). ‘BglBrick vectors and datasheets: a synthetic biology platform for gene expression’. Journal of Biological Engineering 5(1): 1–14.

L. F. Liu & J. C. Wang (1987). ‘Supercoiling of the DNA template during transcription’. Proceedings of the National Academy of Sciences 84(20): 702–7027.

R. Lutz & H. Bujard (1997). ‘Independent and tight regulation of transcriptional units in Escherichia coli via the LacR/O, the TetR/O and AraC/I1-I2 regulatory elements’. Nucleic Acids Research 25(6): 1203–1210.

T. Maier, et al. (2011). ‘Quantification of mRNA and protein and integration with protein turnover in a bacterium’. Molecular Systems Biology 7(1): 511.

S. Meyer, et al. (2014). ‘Torsion-Mediated Interaction between Adjacent Genes’. PLoS Comput Biol 10(9): e1003785.

D. Mishra, et al. (2014). ‘A load driver device for engineering modularity in biological networks’. Nature Biotechnology 32(12): 1268–1275.

T. S. Moon, et al. (2012). ‘Genetic programs constructed from layered logic gates in single cells’. Nature 491(7423): 249–253.

V. K. Mutalik, et al. (2013a). ‘Precise and reliable gene expression via standard transcription and translation initiation elements’. Nature Methods 10(4): 354–360.

V. K. Mutalik, et al. (2013b). ‘Quantitative estimation of activity and quality for collections of functional genetic elements’. Nature methods 10(4): 347–353.

H. C. Nelson & R. T. Sauer (1985). ‘Lambda repressor mutations that increase the affinity and specificity of operator binding’. Cell 42(2): 549–558.

H. Niederholtmeyer, et al. (2015). ‘Rapid cell-free forward engineering of novel genetic ring oscillators’. eLife p. e09771.

V. Noireaux, et al. (2003). ‘Principles of cell-free genetic circuit assembly’. Proceedings of the National Academy of Sciences 100(22): 12672–12677.

M. L. Opel & G. Hatfield (2001). ‘DNA supercoiling-dependent transcriptional coupling between the divergently transcribed promoters of the ilvYC operon of Escherichia coli is proportional to promoter strengths and transcript lengths’. Molecular Microbiology 39(1): 191–198.

Z.-A. Ouafa, et al. (2012). ‘The nucleoid-associated proteins H-NS and FIS modulate the DNA supercoiling response of the pel genes, the major virulence factors in the plant pathogen bacterium Dickeya dadantii’. Nucleic Acids Research 40(10): 4306–4319.

J. S. Paige, et al. (2011). ‘RNA mimics of green fluorescent protein’. Science 333(6042): 642–646.

A. R. Rahmouni & R. D. Wells (1992). ‘Direct evidence for the effect of transcription on local DNA supercoiling *in vivo*’. Journal of Molecular Biology 223(1): 131–144.

K. Y. Rhee, et al. (1999). ‘Transcriptional coupling between the divergent promoters of a prototypic LysR-type regulatory system, the ilvYC operon of Escherichia coli’. Proceedings of the National Academy of Sciences 96(25): 1429–14299.

K. E. Shearwin, et al. (2005). ‘Transcriptional interference-a crash course’. TRENDS in Genetics 21(6): 339–345.

J. Shin & V. Noireaux (2012). ‘An E. coli cell-free expression toolbox: application to synthetic gene circuits and artificial cells’. ACS Synthetic Biology 1(1): 29–41.

D. Siegal-Gaskins, et al. (2014). ‘Gene circuit performance characterization and resource usage in a cell-free breadboard’. ACS Synthetic Biology 3(6): 416–425.

M. J. Smanski, et al. (2014). ‘Functional optimization of gene clusters by combinatorial design and assembly’. Nature biotechnology.

B. C. Stanton, et al. (2014). ‘Genomic mining of prokaryotic repressors for orthogonal logic gates’. Nature Chemical Biology 10(2): 99–105.

J. Stricker, et al. (2008). ‘A fast, robust and tunable synthetic gene oscillator’. Nature 456(7221): 516–519.

Z. Z. Sun, et al. (2013). ‘Linear DNA for rapid prototyping of synthetic biological circuits in an Escherichia coli based TX-TL cell-free system’. ACS Synthetic Biology.

Z. Tuza, et al. (2013). ‘An *in silico* modeling toolbox for rapid prototyping of circuits in a biomolecular “breadboardÓ system’. In Decision and Control (CDC), 2013 IEEE 52nd Annual Conference on, pp. 1404–1410. IEEE.

J.-W. Veening, et al. (2004). ‘Visualization of differential gene expression by improved cyan fluorescent protein and yellow fluorescent protein production in Bacillus subtilis’. Applied and environmental microbiology 70(11): 6809–6815.

E. Weber, et al. (2011). ‘A Modular cloning system for standardized assembly of multigene constructs’. PLoS ONE 6(2).

X. S. Xie, et al. (2008). ‘Single-molecule approach to molecular biology in living bacterial cells’. Annual Reviews of Biophysics 37: 417–444.

J. W. Young, et al. (2012). ‘Measuring single-cell gene expression dynamics in bacteria using fluorescence time-lapse microscopy’. Nature Protocols 7(1): 80–88.

J. Zhang, et al. (2002). ‘Creating new fluorescent probes for cell biology’. Nature Reviews Molecular Cell Biology 3(12): 906–918.

